# Transcriptomic analyses unveil hydrocarbon degradation mechanisms in a novel polar *Rhodococcus* sp. strain R1B_2T from a high Arctic intertidal zone exposed to ultra-low sulfur fuel oil

**DOI:** 10.1101/2025.07.12.664532

**Authors:** Nastasia J. Freyria, Antoine-Olivier Lirette, Brady R.W. O’Connor, Charles W. Greer, Lyle G. Whyte

## Abstract

As Arctic shipping increases due to climate change, characterized by rising temperature and decreasing sea-ice coverage, the risk of oil spills through the Northwest Passage in this fragile ecosystem grows, necessitating effective bioremediation strategies. Research on bioremediation using Arctic coastal sediment bacteria has gained attention, particularly *Rhodococcus* species that play key roles in hydrocarbon degradation (HD) under extreme conditions. This study investigates the HD capabilities of a novel cryophilic Arctic *Rhodococcus* sp. strain R1B_2T isolated from Canadian high Arctic beach sediment in Resolute Bay, exposed to ultra-low sulfur fuel oil for three months at 5 °C. Comparative transcriptomics analyses revealed dynamic responses and metabolic plasticity, with upregulation of genes for aliphatic, aromatic, and polycyclic aromatic hydrocarbons, biosurfactant production (rhamnolipid), cold adaptation, and stress responses. The strain possesses several key alkane degradation genes (*alkB, almA, CYP153, ladA*), with co-expression network analysis highlighting synergistic mechanisms between *alkB* and *CYP153* that target different chain-length alkanes (*alkB*: ∼C5-C20; *CYP153*: ∼C5-C12 and >C30) demonstrating complementary degradation strategies. The findings reveal adaptive mechanisms and degradation kinetics of native Arctic bacteria, highlighting the potential of Arctic cryophilic and halotolerant *Rhodococcus* species for oil spill remediation in polar marine environments.

**HIGHLIGHTS:**

- Novel cryophilic Arctic *Rhodococcus* strain R1B_2T demonstrates robust hydrocarbon degradation capabilities at 4°C
- Transcriptomic analysis reveals upregulation of key genes for aliphatic, aromatic, and PAH degradation pathways
- Synergistic expression between *alkB*, *almA* and *CYP153* genes enables efficient degradation of varied hydrocarbon chain lengths
- Concurrent upregulation of biosurfactant production linked to hydrocarbon biodegradation, cold adaptation, and stress response mechanisms

**GRAPHICAL ABSTRACT:** 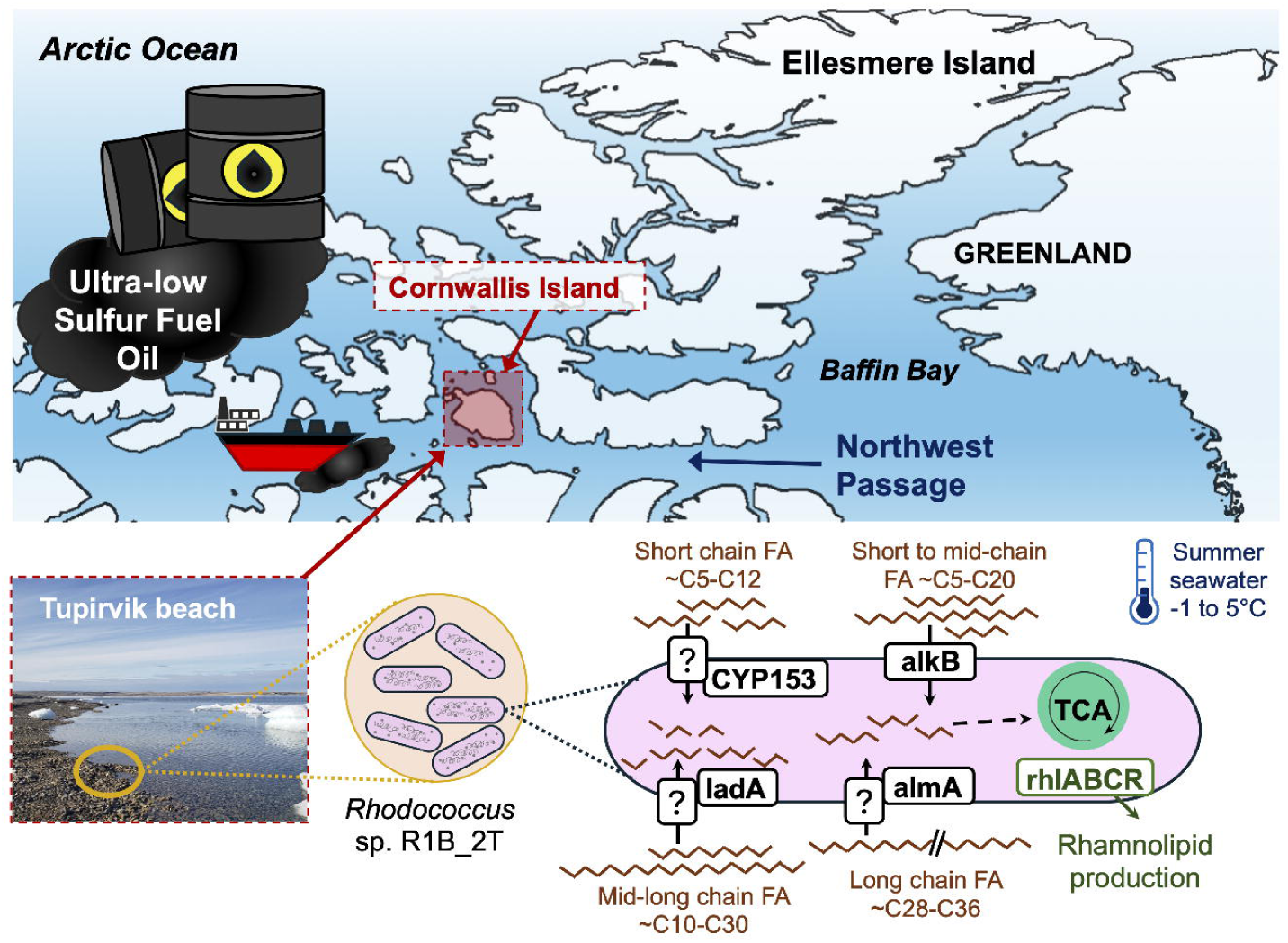

## INTRODUCTION

The Arctic region is experiencing rapid environmental changes with rising temperatures and declining sea-ice thickness and coverage, leading to increased Northwest Passage (NWP) accessibility for maritime traffic (Dawson et al., 2018). This development raises concerns about the heightened risk of oil spills in this pristine and fragile ecosystem (Nevalainen et al., 2017). Heavy fuel oil (HFO) spills pose particular challenges in Arctic conditions due to cleanup difficulties and higher pollutant concentrations (Elbaz et al., 2015; Reddy et al., 2018). While the International Maritime Organization has responded with regulations on HFO use in Arctic waters (Comer et al., 2020), effective bioremediation strategies remain crucial for this environment. Indigenous communities have advocated for stronger protections of their traditional territories (Prior and Walsh, 2018; Dawson et al., 2020), highlighting the need for both improved governance and practical remediation approaches that can function in cold Arctic conditions. Our study addresses this need by investigating the hydrocarbon degradation capabilities of a novel Arctic *Rhodococcus* strain isolated from Canadian high Arctic sediments.

Given the increasing risk of oil spills in the Arctic and the limitations of conventional remediation approaches in this extreme and remote environment, research into the hydrocarbon degradation (HD) capabilities of Arctic coastal sediment bacteria has become paramount for developing effective bioremediation strategies. Arctic coastal sediments harbor diverse microbial communities adapted to extreme cold and fluctuating environmental conditions (Müller et al., 2018), potentially offering native biological solutions for oil contamination that could complement or enhance traditional cleanup methods. These indigenous microorganisms may possess unique metabolic capabilities that enable effective HD under conditions where mechanical recovery or chemical dispersants face important logistical challenges.

Recent studies have identified several Arctic bacterial species with hydrocarbon-degrading capabilities: *Oleispira antarctica* RB-8, which has been identified as an important alkane degrader in cold marine environments (Gregson et al., 2020); *Rhodococcus qingshengii* TUHH-12, which has demonstrated hydrocarbon degradation capabilities in cold conditions (Lincoln et al., 2015); *Glaciecola psychrophila* 170T, a psychrophilic marine bacterium which utilizes cold-active enzymes for aromatic hydrocarbon degradation (Zhang et al., 2006); *Cycloclasticus* sp. 78-ME, which has been studied for aromatic hydrocarbon degradation in cold marine environments (Messina et al., 2016); *Marinobacter* sp. BSs20148, a cold-adapted hydrocarbon degrader from Arctic seawater (Song et al., 2013); and *Polaromonas naphthalenivorans* CJ2, which has been characterized for its ability to degrade naphthalene at low temperatures (Jeon et al., 2006; Jin et al., 2016). These bacteria demonstrate diverse abilities to degrade hydrocarbons in cold environments, highlighting their potential for bioremediation applications. Despite these advances, important knowledge gaps remain regarding the specific metabolic pathways and adaptive mechanisms that enable efficient hydrocarbon degradation in Arctic conditions, particularly for complex fuel mixtures like Ultra-Low Sulfur Fuel Oil (ULSFO). Most studies have focused on single hydrocarbon compounds rather than complex mixtures, and the genetic regulation of cold-adapted degradation pathways remains poorly understood.

Advancements in high-throughput technologies have revolutionized our ability to investigate microbial communities and their functional capacities in extreme environments (Cowan et al., 2015). Techniques such as metagenomics, transcriptomics, and metabolomics allow for comprehensive analyses of microbial responses to oil exposure, providing insights into the genetic and metabolic adaptations that enable HD in cold environments (Kimes et al., 2013; Freyria et al., 2024a; Lirette et al., 2024). Comparative transcriptomic studies of polar sediment and marine bacteria have provided crucial insights into cold-adapted microorganisms’ mechanisms for bioremediation. For instance, Conte et al. (2018) found upregulation of alkane degradation genes and adaptive responses in *Oleispira antarctica* RB-8, while Laczi et al. (2015) revealed differential expression of hydrocarbon metabolism and stress response genes in *Rhodococcus erythropolis* PR4. Qin et al. (2014) demonstrated diverse hydrocarbon-degrading capabilities in the Antarctic bacterium *Glaciecola punicea* ACAM 611T, and Messina et al. (2016) identified and observed upregulation of aromatic HD genes in Arctic *Cycloclasticus* sp. 78-ME. In a more recent study, Gregson et al. (2020) employed comparative transcriptomics to investigate the response of a consortium of Arctic marine bacteria to crude oil exposure. Their research identified key functional genes and metabolic pathways involved in HD, as well as genes related to biosurfactant production and cold adaptation. The Gregson et al. (2020) study emphasized the importance of microbial interactions in effective oil degradation within cold environments. These organisms may possess unique enzymes and metabolic pathways optimized for low-temperature degradation of complex hydrocarbons found in ULSFO (Vergeynst et al., 2019). Understanding their degradation mechanisms could lead to the development of more effective bioremediation strategies specifically designed for Arctic conditions.

Long-term exposure studies in Arctic environments present numerous challenges, including extreme weather conditions, logistical difficulties, and environmental sensitivities (Vergeynst et al., 2018). Despite these obstacles, such studies are crucial for understanding and protecting these fragile ecosystems (Vincent et al., 2011). A notable example includes the Baffin Island Oil Spill (BIOS) study (Owens et al., 1987; Sergy and Blackall, 1987; Hunnie et al., 2023; Schreiber et al., 2023), which has provided valuable insights into the fate and effects of oil in Arctic marine ecosystems over four decades. By combining in-situ observations with laboratory experiments and high-throughput analytical techniques, researchers can gain a comprehensive understanding of the oil degradation capacities of Arctic marine bacteria. This knowledge is essential for developing predictive models and informed strategies for oil spill response in the Arctic, where traditional clean-up methods may be less effective due to the remote location and harsh environmental conditions (Vergeynst et al., 2018).

Our study focuses on *Rhodococcus* sp. strain R1B_2T, a psychrotolerant bacterium with growth capacity ranging from 0 °C to 25 °C, isolated from high Arctic NWP beach sediment, Resolute Bay, Nunavut, Canada (Lirette et al., 2024). This adaptation to cold environments makes it particularly relevant for studying hydrocarbon degradation in Arctic conditions. In our recent metagenomic, and culture-based investigations of NWP marine beach sediments (Freyria et al., 2024a; Góngora et al., 2024; Lirette et al., 2024), *Rhodococcus* appears to play a predominant role in HD (Whyte et al., 1998; Whyte et al., 2002) and is considered one of the dominant genera in oil-contaminated Arctic marine beach sediments (Whyte et al., 2002), highlighting its potential for in-situ bioremediation (Whyte et al., 2001). The objective was to ascertain the potential of *Rhodococcus* strain R1B_2T to degrade aliphatic, aromatic and polycyclic aromatic hydrocarbons, and to identify the key genes and key metabolic pathways involved. This was achieved by exposing the strain to ULSFO as the sole carbon source for a period of three months at 5 °C in liquid cultures. Comparative transcriptomics analyses were performed to reveal a dynamic response of *Rhodococcus* and demonstrate its metabolic plasticity in cold marine environments.

## EXPERIMENTAL PROCEDURES

### Strain isolation and growth conditions

Lirette (2023) isolated *Rhodococcus* strain R1B_2T. Briefly, sediments (5 g) from a 1-month column experiment simulating the intertidal cycle in Arctic beach sediment (Chen et al., 2024) were placed in 50 mL Falcon tubes, which were then supplemented with sterilized glass beads and water (3 times the sediment weight, 1:4 dilution). Each tube was vortexed for 2 minutes to dislodge the sediments. Subsequently, 1 mL of each solution was added to 9 mL of autoclave distilled water, achieving a 10-fold dilution. This process continued until a 10^-5^ dilution was reached. Then, 100 μL of supernatant from the 10^-3^, 10^-4^, and 10^-5^ dilutions were spread on two types of plates: Reasoner’s 2A (R2A, BD) gellan plates (Alfa Aesar), which support slow-growing microorganisms (Reasoner and Geldreich, 1985), and plates made of autoclaved beach water (from Tupirvik beach, Resolute Bay), 2% gellan gum (Alfa Aesar), and 0.67 g.L^-1^ ammonium sulfate amended with 500 ppm ULSFO (Lirette, 2023).

To determine hydrocarbon biodegradation, ULSFO was added to a final concentration of 500 ppm (0.045 g of ULSFO per 100 mL) of sterile artificial sea water (3% sea salt), with triplicates for 3 conditions: i) negative control culture without *Rhodococcus* cells and supplemented with 500 ppm of ULSFO and 0.35 g.L^-1^ ammonium sulfate; ii) control culture of *Rhodococcus* cells without ULSFO and supplemented with R2A broth and 0.35 g.L^-1^ ammonium sulfate; iii) and iv) culture of *Rhodococcus* cells supplemented with 500 ppm of ULSFO and 0.35 g.L^-1^ ammonium sulfate. Cultures of *Rhodococcus* under all conditions were incubated for 1 and 3 months on shakers at 100 rpm in a 5 °C incubator (Lirette, 2023). The initial concentration of cells was 10^4^ cells.mL^-1^. Cells were collected after each time point (T0 = initial time point, T1 = after 1 month, and T3 = after 3 months) in 50 mL falcon tubes, centrifuged for 2 min at 5,000 rpm, and supernatant was removed each time. Pellets were resuspended in 2 mL of Zymo Research DNA/RNA Shield (Irvine, CA, USA), incubated for 30 min at room temperature and stored at −80 °C until processed.

### Petroleum hydrocarbon analyses

After each incubation period, the medium was heated to 60 °C to suspend ULSFO adhering to the glass flask, then transferred to glass bottles provided by SGS Canada (Montreal, Quebec) and stored at −20 °C until all samples were collected for petroleum hydrocarbon analysis. For aqueous samples, the extraction of semi-volatile organic compounds (SVOCs) was carried out using dichloromethane by liquid-liquid extraction to ensure complete removal of the analytes from the water, and the quantification of SVOCs was conducted using the United States Environmental Protection Agency (EPA) Method 8270E (EPA, 2014). Petroleum hydrocarbons, including polycyclic aromatic hydrocarbons (PAHs) and aliphatic hydrocarbons, were analyzed according to SGS Canada’s protocols and the CCME Reference Method for Petroleum Hydrocarbons in Soil – Tier 1 (Canadian Council of Ministers of the Environment (CCME), 2002). The initial concentration of each hydrocarbon group in ULSFO was determined by employing negative controls of culture without cells as they demonstrated no indications of degradation. The hydrocarbon removal rate was determined by subtracting the measured amounts of each group in the remaining oil and liquid culture from the initial concentration in ULSFO (**Table S1**). This result was divided by the concentration of that group in ULSFO and multiplied by 100 to obtain the percentage of hydrocarbon removal (Chen et al., 2024). All experimental samples were analysed with biological triplicates (three independent cultures), each with technical duplicates (n = 6 total measurements per condition). The negative control (medium with oil but no cells, T0) was analyzed with technical duplicates only. Errors bars represent the standard deviation across all replicates for each condition.

### Whole genome sequencing and processing

Whole genome extraction was conducted using the DNeasy PowerSoil DNA extraction kit (Qiagen) according to the manufacturer’s instructions. Whole genome sequencing was performed with DNA shotgun (with PCR) following the sample preparation guidelines: 50 µL of solution containing 150 ng of nucleic acid was added to a 96-well plate. Library preparation was performed using the NEBNext Ultra II DNA Library Prep Kit for Illumina according to the manufacturer’s protocol. DNA concentrations were verified with a Qubit fluorometer (Life Technologies, Invitrogen) using the dsDNA Kit (Invitrogen) and with a NanoDrop 8000 spectrophotometer (Thermo Scientific). The sequencing was performed at Genome Quebec (Montréal, Canada) using the Illumina NovaSeq 6000 S4 PE150 platform, resulting in the generation of 35M reads. Raw reads were trimmed using Trimmomatic v0.36 (Bolger et al., 2014) and assembled with Spades v3.15.4 (Bankevich et al., 2012). Assembled reads were annotated using METAerg (Dong and Strous, 2019) for Kyoto Encylopedia of Genes and Genomes (KEGG) pathways (Kanehisa and Goto, 2000). The Calgary approach to annotating hydrocarbon (CANT-HYD) (Khot et al., 2022) was used to detect hydrocarbon degradation genes with an E-value cut-off of 0.01. Assembled reads were also submitted to JGI for inclusion in the Integrated Microbial Genomes (IMG) database and functional annotation (Markowitz et al., 2007).

### RNA extraction, library preparation and sequencing

RNA from cells were extracted using RNeasy mini kit (Qiagen) according to the manufacturer’s instructions. Quantification of RNA was verified using a Qubit fluorometer using the RNA BR Assay Kit and the quality was checked with NanoDrop 8000 spectrophotometer. Sequencing libraries were prepared with the NEBNext rRNA Depletion kit (Bacteria, New England BioLabs). The DNA libraries were then pooled equimolarly for normalization. The quality and quantity of the pooled libraries were verified using the Agilent High Sensitivity DNA kit on a 2100 Bioanalyzer (Agilent Technologies, Santa Clara, CA, USA). The indexed and pooled final library was sequenced on an Illumina NexSeq llumina sequencer with PE100 flow cell at Genome Quebec Innovation Center, resulting in the generation of 400M reads.

### Sequence data processing

The experiment involved 15 samples, derived from three different time points (T0, T1, and T3), each with three replicates. These samples were split across three conditions: the control group with cells and without ULSFO at T0, T1, and T3, and cells exposed to ULSFO at T1 and T3. The following analyses were conducted using Compute Canada (Digital Research Alliance) facilities and in-house computers. Raw sequences were quality checked at Genome Quebec and were trimmed in-house using BBMap v.38.18 (Bushnell, 2014) with paired-end mode. All trimmed transcriptome reads were mapped separately onto the *Rhodococcus* strain R1B_2T reference genome (Lirette et al., 2024) available at the JGI (Joint Genome Institute) Genome Portal using Spliced Transcripts Alignment to a Reference (STAR) v.2.7.10b (Dobin and Gingeras, 2015) with default parameters. The mapped reads were counted to evaluate the number of aligned read pairs to each gene between biological replicates. Overall, transcriptome reads mapped to 4,395 genes in the *Rhodococcus* strain R1B_2T reference genome. The total number of clean reads generated for all samples was 697,108. After the number of reads per gene were determined for each transcriptome, sequencing replicates were pooled (**Table S2**).

Gene annotations were retrieved from the JGI Genome Portal with *Rhodococcus* strain R1B_2T as the reference genome. Gene annotations included those from Gene Ontology (GO), KEGG, eukaryotic Ortholog Groups of proteins (KO), Clusters of Orthologous Genes (COG), signal peptide and Interproscan annotations. Carbohydrate active enzymes (CAZyme) were predicted using dbCAN, a web server for automated CAZyme annotation using hmmscan v.3.3.2 and based on the CAZy database (Cantarel et al., 2009). The CANT-HYD (Khot et al., 2022) was utilized to detect the presence of marker genes among all transcriptome reads, with a ‘noise’ cut-off score. Hydrocarbon Aerobic Degradation Enzymes and Genes (HADEG) (Rojas-Vargas et al., 2023) was also used to detect maker genes. Both CANT-HYD and HADEG enables the identification of genes involved in both aerobic and anaerobic hydrocarbon degradation pathways, encompassing aliphatic and aromatic hydrocarbons.

### Phylogeny and pangenome comparison

A phylogenetic tree was constructed as previously described (Freyria, 2021; Freyria et al., 2021) from curated full-length 16S rRNA gene sequences of the several known species of *Rhodococcus* using randomized accelerated maximum likelihood (RAxML) v.8.2.11, 1000 bootstrap replicates and using the model GTRGAMMA (Stamatakis, 2014). Sequences were aligned using MUSCLE (v.3.8.31) (Edgar, 2004). The genome of *R.* sp. strain R1B_2T (Lirette, 2023) was compared to the genome of *R. cercidiphylli* strain IEGM 1322 (RefSeq: GCF_033042225.1) using OrthoANI calculations from EzBioCloud (Chalita et al., 2024). The average nucleotide identity (ANI) analysis was performed using fastANI v.1.34 (Hernández-Salmerón and Moreno-Hagelsieb, 2022) by comparing 8 genomes of *Rhodococcus* species and the genome of *Rhodococcus* sp. strain R1B_2T (**Table S3**).

Pangenome analysis was performed using Roary v.3.13.0 (Page et al., 2015) by comparing the 9 mentioned genomes, following rapid functional annotation using Prokka v.1.14.6 (Seemann, 2014). To examine the evolutionary relationships of key genes involved in hydrocarbon degradation, we constructed phylogenetic trees for *alkB* (using MUSCLE alignment and RAxML with GTRGAMMA model) and *almA* genes. For *almA,* putative homologs were identified by blastp against the known *almA* gene from *Alloalcanivorax dieselolei* (ADP30851.1; **Table S4**), and a phylogenetic tree was constructed from the highest-scoring sequences using RAxML with model PROGAMMALG and 1,000 bootstrap replicates.

### Statistical analyses

All further analyses were performed in the RStudio v.4.3.3 environment. Counted mapped reads of all transcriptomes were normalized using *DESeq2* package in R (Love et al., 2014), where counts are divided by sample-specific size factors determined by the median ratio of gene counts. This step was followed by differential gene expression analyses using *DESeq2*. Multiple hypothetical testing was corrected by the Benjamini-Hochberg method (Benjamini and Hochberg, 2018) included in the *DESeq2* package. Genes with Benjamini-Hochberg-adjusted *p*-value ≤0.05 and an absolute fold change (FC) ≥2 (log2 FC ≥2) were identified as significant differentially expressed genes (DEGs). These DEGs were observed between the different sampling times (T0, T1, and T3) and between the control (without oil) and the oil incubation conditions. The representation of DEGs between samples was visualized using scatter plots as previously described (Freyria et al., 2022).

A two-way analysis of variance (ANOVA) using Past4 v.4.11 software, but with repeated measure was conducted to analyze the time effect between conditions on TPH analysis. Canonical Correspondence Analysis was computed using the *cca()* function from the *vegan* package (Dixon, 2003) to discriminate different transcriptomes for each condition according to the time of sampling and the effect of the presence of ULSFO. A dendrogram was computed using the *hclust*() function from the *Stats* package (R Core Team, 2018). The complete linkage method was used to construct the dendrogram, which finds similar clusters among samples. The top 25 DEG heatmap was computed on normalized and transformed read counts with the *vst()* function in *Deseq2* and visualized with the *pheatmap()* function from the *pheatmap* package (Kolde and Kolde, 2015). Circular heatmaps of selected DEGs were based on the normalized number of reads per gene and on values of log_2_ FC between comparison of time of sampling among all conditions, and were visualized using the *circos.heatmap()* function in R from the *circlize* package (Gu et al., 2014). To visualize the DEGs with differential gene expression relative to the log-fold change, we used scatter plots and volcano plots computed with the *DESeq2* package.

We performed Weighted Gene Co-expression Network Analysis (WGCNA) using *WGCNA* package (Langfelder and Horvath, 2008) in R on DEGs between conditions to analyze conditions of incubation of ULSFO and time effects as previously described (Freyria et al., 2024b). To construct co-expression modules, we used default settings except with a soft thresholding power of 15, a minimum Module size of 50 genes, and a branch merge cut height of 0.25. Genes were clustered into 4 correlated modules. To identify the relationship between the time of sampling and oil conditions within all modules, the Pearson’s correlation coefficient was calculated. Hub genes with the highest connectivity among each module were calculated using intramodular connectivity (K.in) and module correlation degree (MM) of each gene. Hub genes may represent key genes with potentially important functions. Module Membership (MM) vs Gene Significance (GS) was statistically investigated in four modules (brown, blue, turquoise and grey) via the WGCNA package. MM refers to the strength of the association between a gene and a module. GS is defined as the correlation between gene expression and stress status. Networks of each module were visualized using Cytoscape v.3.9.1 (Shannon et al., 2003). The node circle size is positively correlated with the number of genes that are partnered within interactions.

## RESULTS

### Hydrocarbon biodegradation analyses

Analysis of residual fuel after one and three months (T1 and T3) of exposure to *Rhodococcus* strain R1B_2T showed similar percentages of ULSFO removal due to biodegradation at both time points (**Fig. 1, Table S1**). For the alkane biodegradation, almost 80% of fraction 2 (C10-C16) was removed after a month (**Fig. 1A**). The alkane fraction shows a clear and significant decrease (one-way ANOVA, *p*-value = 0.00171 **; **Table S5**) between one and three months of fuel exposure for only alkane fraction F3 (C16-C34; **Fig. 1B**). For the aromatic compounds, the results are more varied (**Fig. 1C-1D**). The degradation of 1-Methylnaphthalene, 2-Methylnaphthalene, Methylnaphthalene 2-(1-) and Naphthalene do not show statistically significant changes (one-way ANOVA*, p*-value >0.05; **Table S5**). The percentage of removal of naphthalene was higher than the methylated-naphthalene but by comparing the initial concentration of aromatic compounds, there were lower naphthalene concentrations in ULSFO than the methylated-naphthalene (**Fig. 1D**). Overall, a Wilcoxon test revealed significant difference in ULSFO removal percentages between T1 and T3 (*p*-value >0.05) for alkane fractions, while for aromatic compound, no significant differences were found.

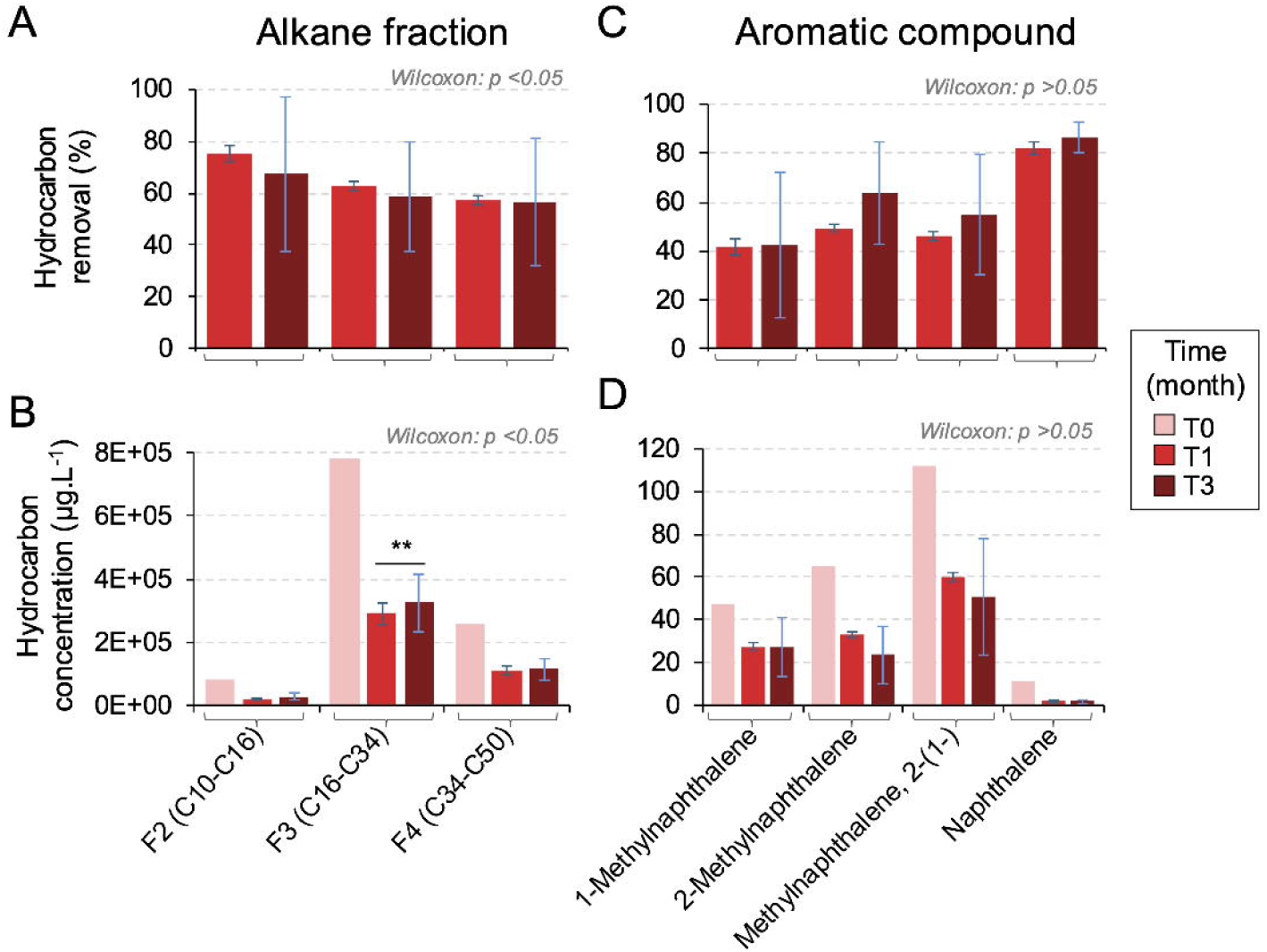

### Novel subspecies of Arctic NWP beach sediment genus *Rhodococcus*

To determine the taxonomic position of *Rhodococcus* strain R1B_2T isolated from Arctic NWP beach sediment, we performed comprehensive phylogenetic and genomic analyses. A maximum likelihood (RAxML) phylogenetic tree based on 16S rRNA gene sequences from 56 known *Rhodococcus* species and strains positioned R1B_2T in close proximity to *R. cercidiphylli* (**Fig. 2A**). The broad cluster containing R1B_2T, *R. cercidiphylli*, and *R. cerastii* is supported by a bootstrap value of 85, indicating good statistical support for this grouping. However, the specific branching pattern between R1B_2T and *R. cercidiphylli* lacks bootstrap support, suggesting some uncertainty in the fine-scale relationship despite their clear affiliation. Notably, R1B_2T clustered distinctly from other known polar *Rhodococcus* species such as *R. erythropolis* and *R. qingshengii*.

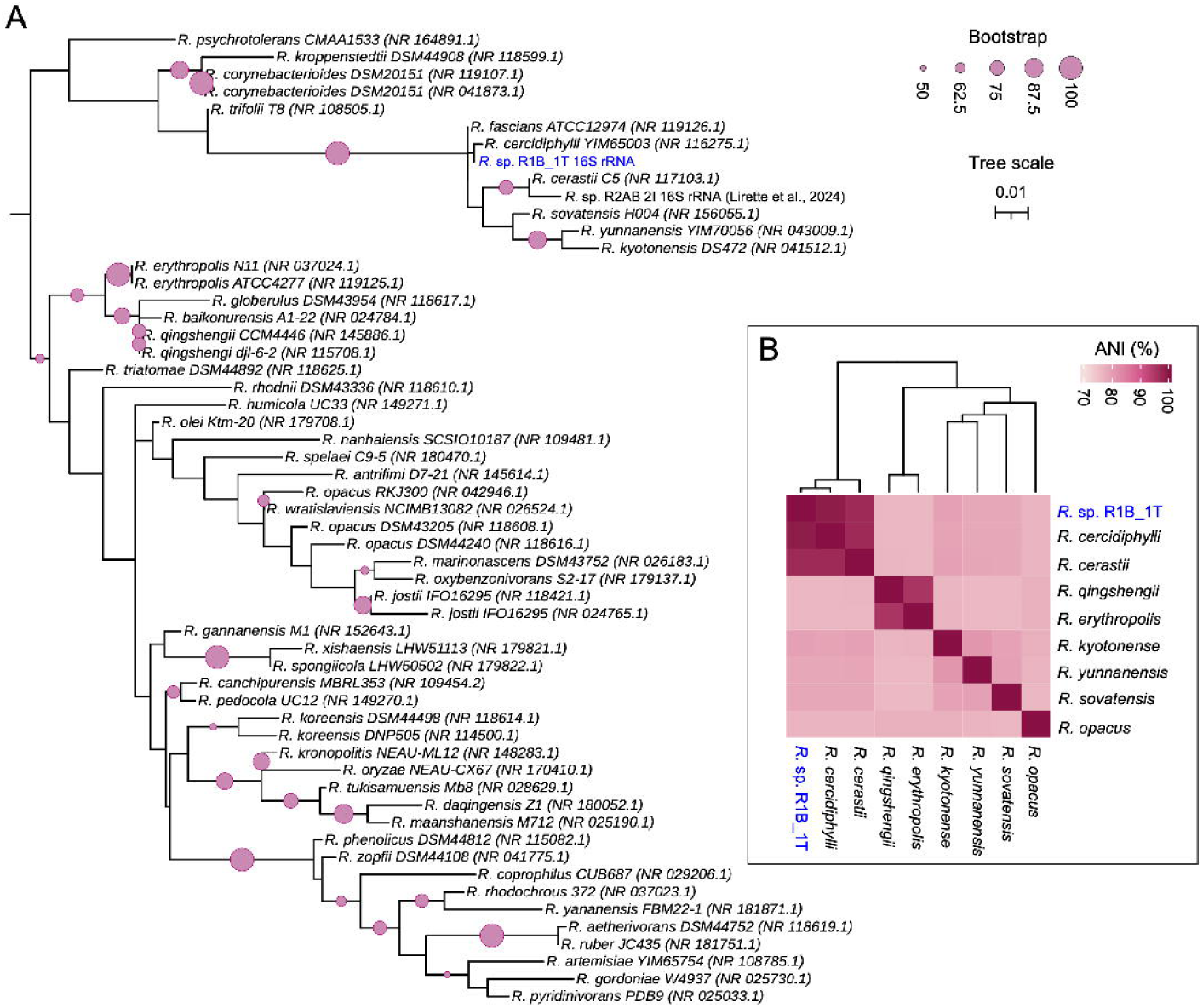

Whole genome comparison using OrthoANI calculations revealed 99% similarity between R1B_2T and *R. cercidiphylli* strain IEGM 1322 (RefSeq: GCF_033042225.1). The genome sizes are nearly identical, with R1B_2T at 5,497,800 bp and *R. cercidiphylli* strain IEGM 1322 at 5,489,640 bp. Detailed Average Nucleotide Identity (ANI) analysis confirmed high genomic similarity between R1B_2T and *R. cercidiphylli* (98.9%) and *R. cerastii* (97.1%), while showing substantial divergence from *R. qingshengii* and *R. erythropolis* (76.27% and 76.24%, respectively; **Fig. 2B, Table S3**).

Pangenome analysis comparing R1B_2T with 8 reference *Rhodococcus* species genomes (**Fig. S1, Table S3**), revealed that only 0.5% of genes constitute the core genome shared across all 9 genomes. The analysis identified 17.4% cloud genes (present in <15% of genomes), 14% shell genes (shared between 15-95% of genomes), and a substantial proportion of unique genes specific to individual genomes. These genomic patterns, combined with the ANI values exceeding 98% with *R. cercidiphylli*, indicate that strain R1B_2T represents a novel subspecies of *R. cercidiphylli* rather than a distinct species, adapted to the unique conditions of Arctic beach sediments.

Phylogenetic analysis of *alkB* and *almA* genes across nine *Rhodococcus* genomes (**Fig. S2**) revealed that strain R1B_2T possesses multiple copies of these genes, clustering distinctly from those found in *R. erythropolis* and *R. qingshengii*. The three *alkB* genes from R1B_2T formed a strongly supported clade (bootstrap value >99%) with *R. cercidiphylli* and *R. cerastii* (**Fig. S2A**), consistent with our previous whole-genome phylogenetic analysis, while showing clear separation from the *alkB* genes of known Arctic hydrocarbon degraders *R. erythropolis* and *R. qingshengii*. Similarly, the *almA*-like flavin-binding monooxygenase from R1B_2T clustered with *R. cercidiphylli* and *R. cerastii*, distinct from those of *R. erythropolis* and *R. qingshengii* (**Fig. S2B, Table S4**).

### An overview of transcriptome assembly and annotation

Differential gene expression from all 15 transcriptomes from all triplicate conditions at all timepoints showed a similar clustering pattern using three methods: a cluster dendrogram (**Fig. 3A**), a constrained correspondence analysis ordination (**Fig. 3B**), and a “top” 25 most variant genes heatmap (**Fig. 3C**). The cluster dendrogram showed distinct grouping of samples, with oil-treated samples at T1 and T3 clustering separately from no-oil control samples. The CCA ordination showed a separation between samples into distinct groups based on their experimental conditions with the control samples forming one cluster, while the oil-treated samples (T1 and T3) formed a separate cluster. The first two principal components explain a substantial portion of the variance (23.92% and 19.03%, respectively). The heatmap displayed the expression levels of the top 25 genes with the highest variance based on normalized and transformed read counts across all samples. The heatmap shows distinct expression patterns between the oil-treated and control samples, with 4 genes in cluster b being upregulated in oil treatment samples, while the remaining 21 genes in clusters a and c (a: 4 genes; c:17 genes) showed higher expression in control samples (**Fig. 3C**). Based on expressed genes, samples separated into 3 vertically arranged gene centric clusters. Among the top 25 most variant genes in ULSFO-exposed samples at T1 and T3, we identified genes associated with fatty acid (FA) degradation, transport and modification, including a cytochrome P450 (also known as *CYP153*) and *fadD* (K01897), a long-chain acyl-CoA synthetase (**Fig. 3C, Table S6**). The rest of the top 25 genes were mostly genes annotated in the ‘Translation’ and ‘Post-translation’ categories.

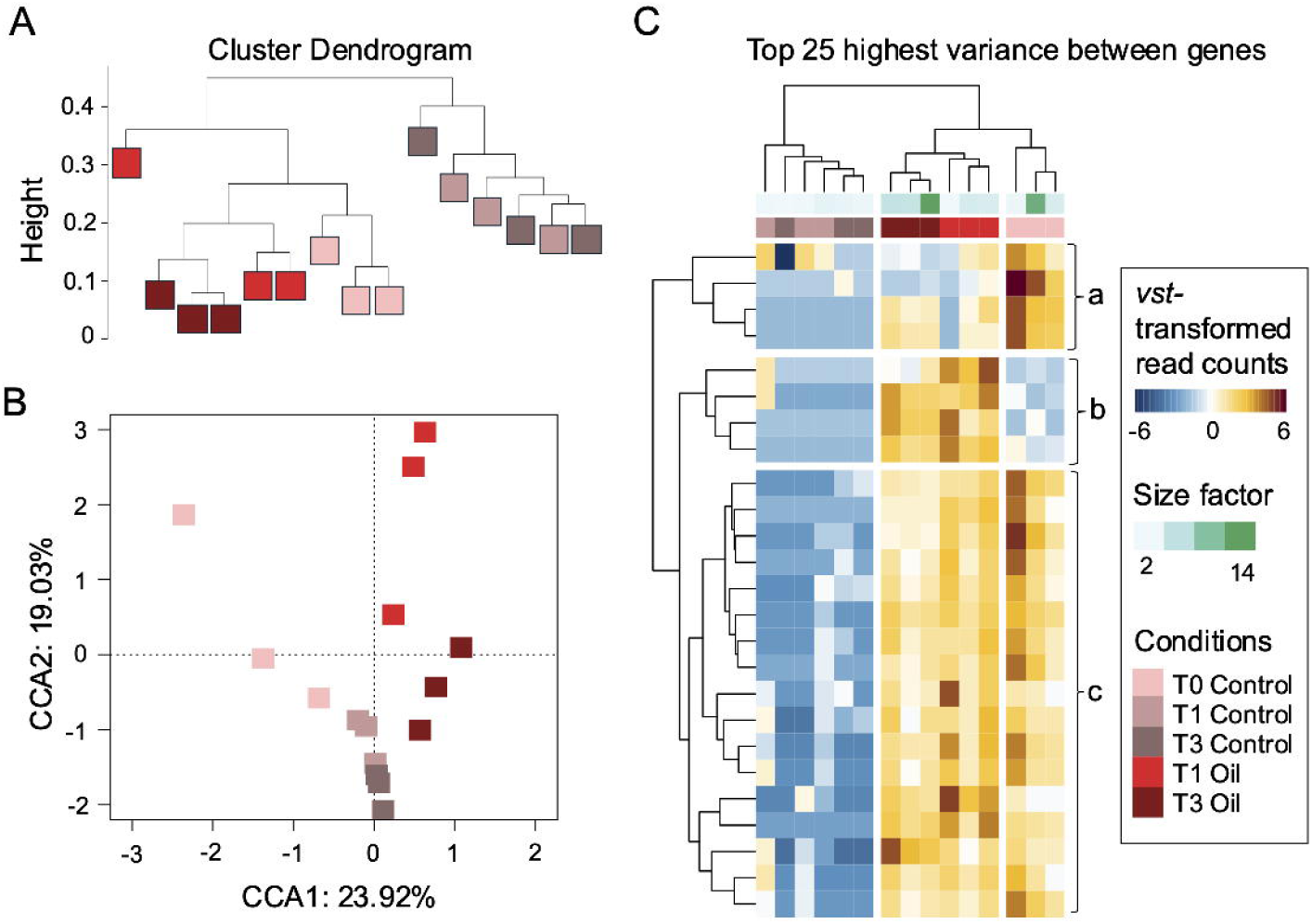

### Differential gene expression over time exposed to oil

Throughout our analyses, we identified differentially expressed genes (DEGs), either up- or down-regulated) over the time of exposure to ULSFO. In comparing the time effect in gene expression, we observed that control comparisons (T0 vs T1 Control, T0 vs T3 Control, T1 vs T3 Control) showed relatively balanced changes in gene expression, with similar numbers of up- and down-regulated genes (**Fig. S3**). The comparison between control and oil treatments at the same time points (T1 Control vs T1 Oil, T3 Control vs T3 Oil) reveals substantial differences, including the highest numbers of DEGs (1,219) at one month exposure (T1 Oil) versus the control at the same time (Control T1). Oil treatments (T0 vs T1 Oil, T0 vs T3 Oil) resulted in more pronounced changes, with the lowest number of DEGs (140) at T3 Oil versus T1 Oil. The dot plots showed the significant DEGs (defined as DEGs with a p-value <0.05), and this stringency reduced the number of DEGs considered (**Fig. S3**).

### Overall functional annotations

To further glean biological meaning from the DEGs resulting from the ULSFO exposure over time, we examined the placement of genes within the GO, COG and KEGG databases (**Figs. 4, S4-S5**). The GO terms provide functional annotation of gene products in three domains: Biological Process (BP; **Fig. 4A**), Cellular Component (CC; **Fig. 4B**), and Molecular Function (MF; **Fig. 4C**). In the BP domain, cells exposed to ULSFO at T1 and T3 had more DEGs annotated within three main processes: 16 genes involved in tricarboxylic acid cycle (TCA; **Table S7**), 11 genes involved in FA beta-oxidation, and 9 genes involved in carbohydrate transport. In the CC domain, a maximum of 168 and 154 DEGs encoded for ‘plasma membrane’ at T1 oil versus T1 control and at T3 oil versus T3 control, respectively (**Fig. 4B, Table S7**). For the same comparison, we observed the highest number of DEGs encoding for ‘cytoplasm’. In the MF domain, a maximum of 153 DEGs were observed at T1 oil versus T1 control within the ‘ATP binding’.

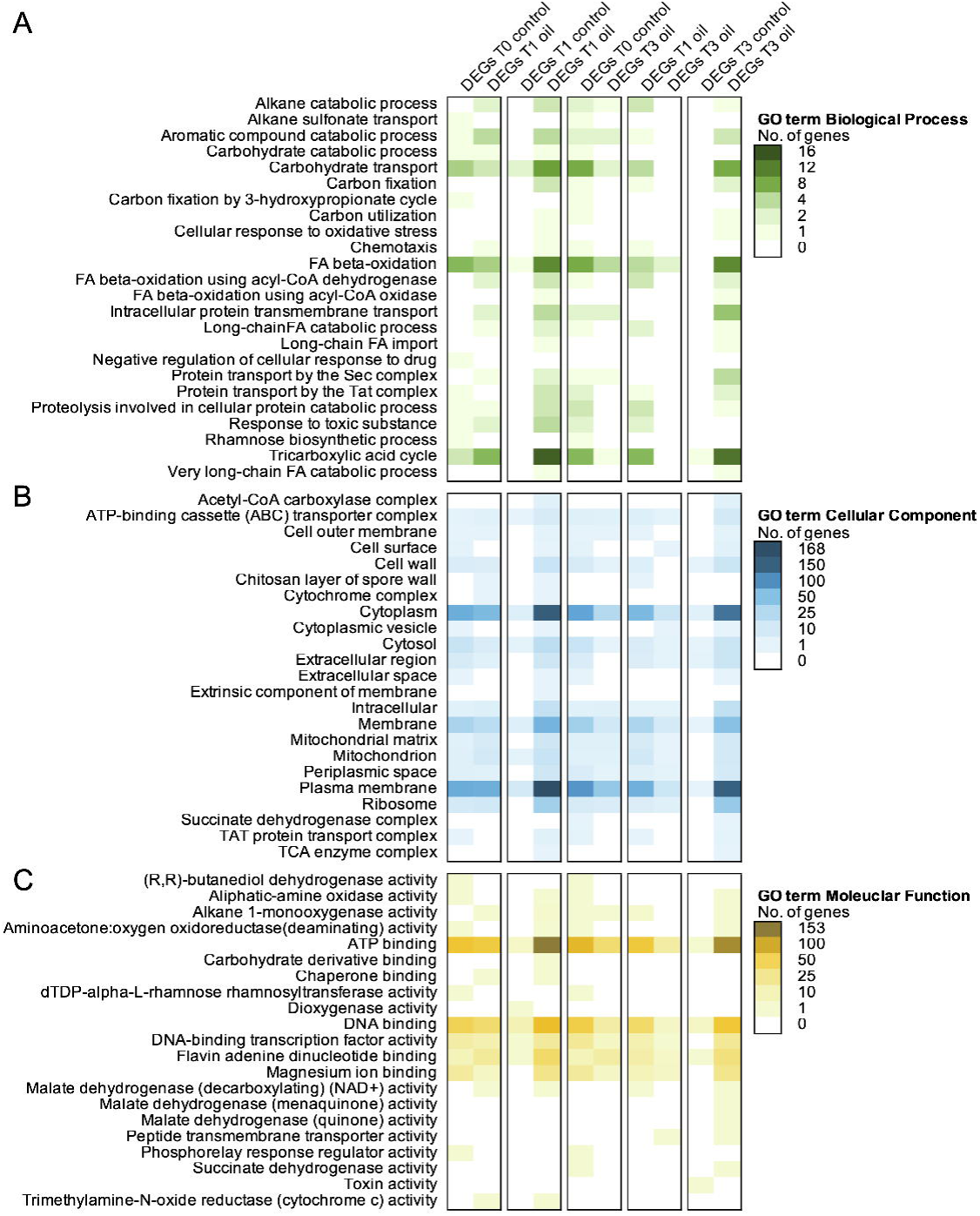

Functional annotation from the COG database revealed two categories, where we observed the highest number of DEGs (5 genes at T1 Oil vs T1 Control) encoding for an acyl-CoA dehydrogenase related to the alkylation response protein AidB (COG1960) involved in FA biosynthesis; and ca. 4 DEGs encoding for an acyl-CoA reductase related to aldehyde dehydrogenase (COG1012) involved in proline degradation (**Fig. S4, Table S8**). At T3 Oil vs T3 Control, we observed the highest number of DEGs (4 genes) within the category of FA desaturase (COG3239) and enoyl-CoA hydratase/carnithine (COG1024) involved in FA biosynthesis. Within the KEGG database, in several pathways, we observed in several pathways the overexpression of genes especially at T1 Oil vs T1 Control, with 63 DEGs involved in FA metabolism (ko01212) and 56 DEGs involved in propanoate metabolism (ko00640; **Fig. S5, Table S9**).

### Co-expression network of differentially expressed genes

We analyzed the expression patterns of 1,037 DEGs identified across all transcriptomes. Using weighted gene co-expression network analysis (WGCNA; **Fig. 5**), the DEGs were categorized into four distinct modules based on their gene expression profiles (**Fig. 5A**). In the cluster dendrogram of eigengene distances, which reflects DEGs grouping patterns, two main branches emerged, with the Brown module distinctly separating from the Turquoise, Blue and Grey modules.

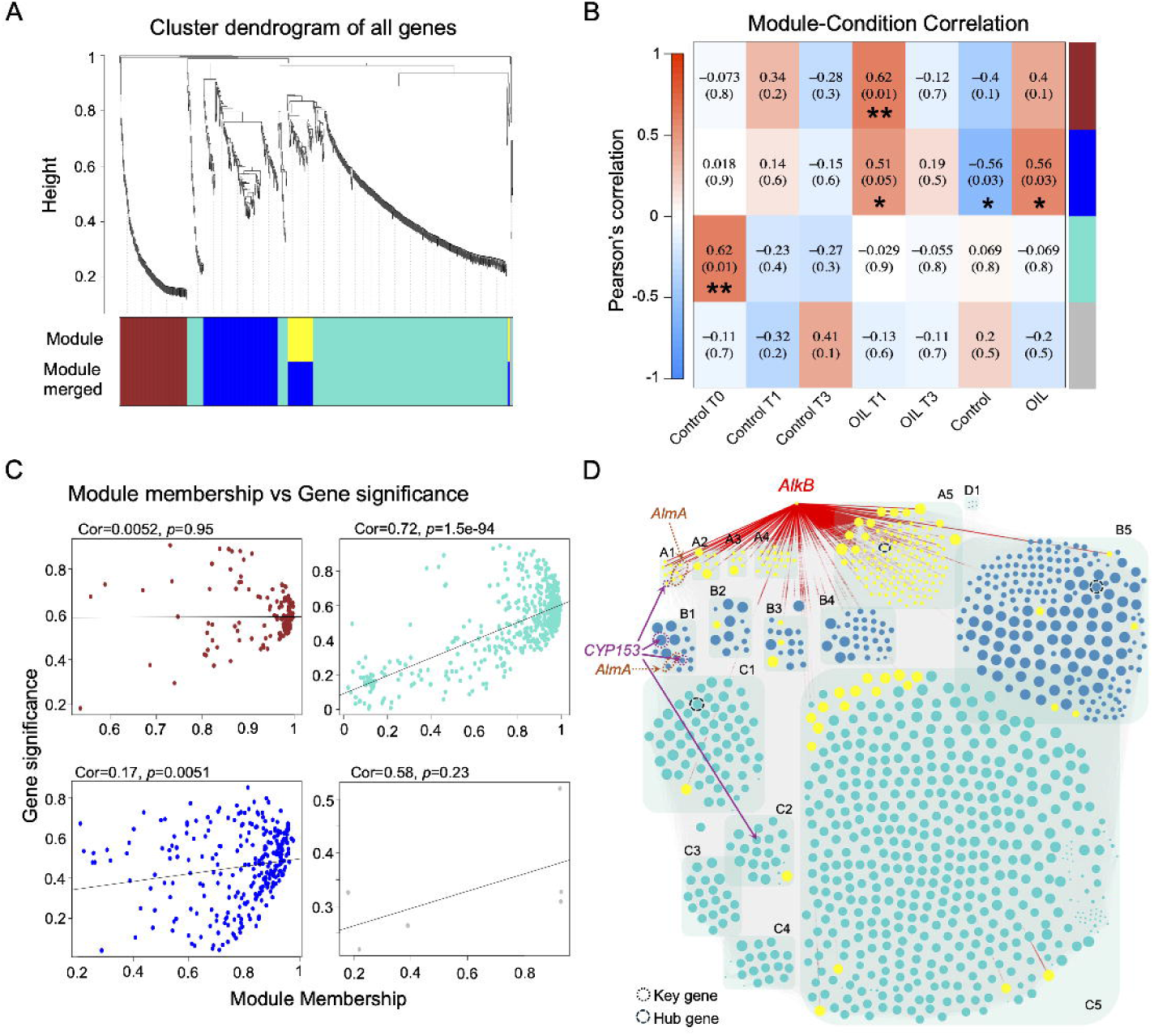

Several modules displayed significant correlations with specific sampling times under either ULSFO exposure or control conditions (**Fig. 5B**). The Blue module showed significant positive correlation with the T1 Oil condition and all oil conditions (both T1 and T3 Oil; *p*-value <0.05 *), while negatively correlating with all control conditions (both T1 and T3 Control; *p*-value <0.05 *). The Brown module was specifically associated with the T1 Oil condition (*p-*value <0.01 **). In contrast, the Turquoise module exhibited significant positive correlation with the T0 Control condition (*p-*value <0.01 **), identifying a gene set that is highly expressed under normal conditions but downregulated during oil exposure. This Turquoise module represents cellular processes that are suppressed when metabolic resources are redirected toward hydrocarbon degradation, complementing the oil-responsive gene networks in the Blue and Brown modules.

The correlation between gene significance and module membership varied substantially across modules (**Fig. 5C**). The Turquoise module displayed the strongest correlation (Cor = 0.72, *p*- value = 1.5e-94), indicating genes central to this module were highly associated with the normal condition and systematically downregulated during oil exposure. The Blue module showed a weaker but significant correlation (Cor = 0.17, *p*-value = 0.0051), while the Brown module, despite its strong association with oil exposure exhibited no correlation (Cor = 0.0052, *p*-value = 0.95), suggesting its response involves complex regulation where connectivity does not predict responsiveness. These patterns reveal distinct organizational principles in gene networks responding to hydrocarbon stress.

To investigate functional relationships between key hydrocarbon degradation genes, we constructed a focused network highlighting connections of *alkB*, *almA*, and *CYP153* genes within our co-expression modules (**Fig. 5D**). Our analysis encompassed 1,037 DEGs (nodes) with 263,410 connections (edges) across four modules. The alkane 1-monooxygenase (*alkB*) gene demonstrated extensive connectivity with all DEGs in the Brown module, which strongly correlated with oil exposure conditions (**Fig. 5A-C**), while also forming connections with multiple DEGs in both Blue and Turquoise modules. Notably, the *alkB* gene from the Brown module (upregulated under ULSFO exposure) selectively connected with *CYP153* and *almA* genes within the same module, but not with their counterparts in the Blue and Turquoise modules, suggesting module-specific coordination of hydrocarbon degradation pathways. Each module featured distinct hub genes: the Blue module’s hub was a choline/glycine/proline betaine transport protein (*proV*, K02168) with 66 nodes and 790 connections; the Brown module’s hub was glycosyltransferase 4 (GT4), involved in cell wall biosynthesis, with 56 nodes and 694 connections; and the Turquoise module’s hub was glycosyl hydrolase 13 subfamily 33 (GH13_33), functioning as a trehalose synthase and/or maltose glucosylmutase.

### Differentially expressed genes of specific pathways

We selected only DEGs that were identified to a specific annotation, or potential pathways involved in HD genes. This was achieved by leveraging the findings of module networks (**Fig. 5**) and investigating functional annotations (**Figs. 4, 6, S4-S7**). A schematic representation of the *Rhodococcus* cell is provided, offering a summary of the potential roles and implications of the selected DEGs in the response of the *Rhodococcus* exposed to ULSFO over a three-month period (**Figs. 7-8**). Several categories of pathways and key genes from **Figs. 7 to 8** are further developed below.

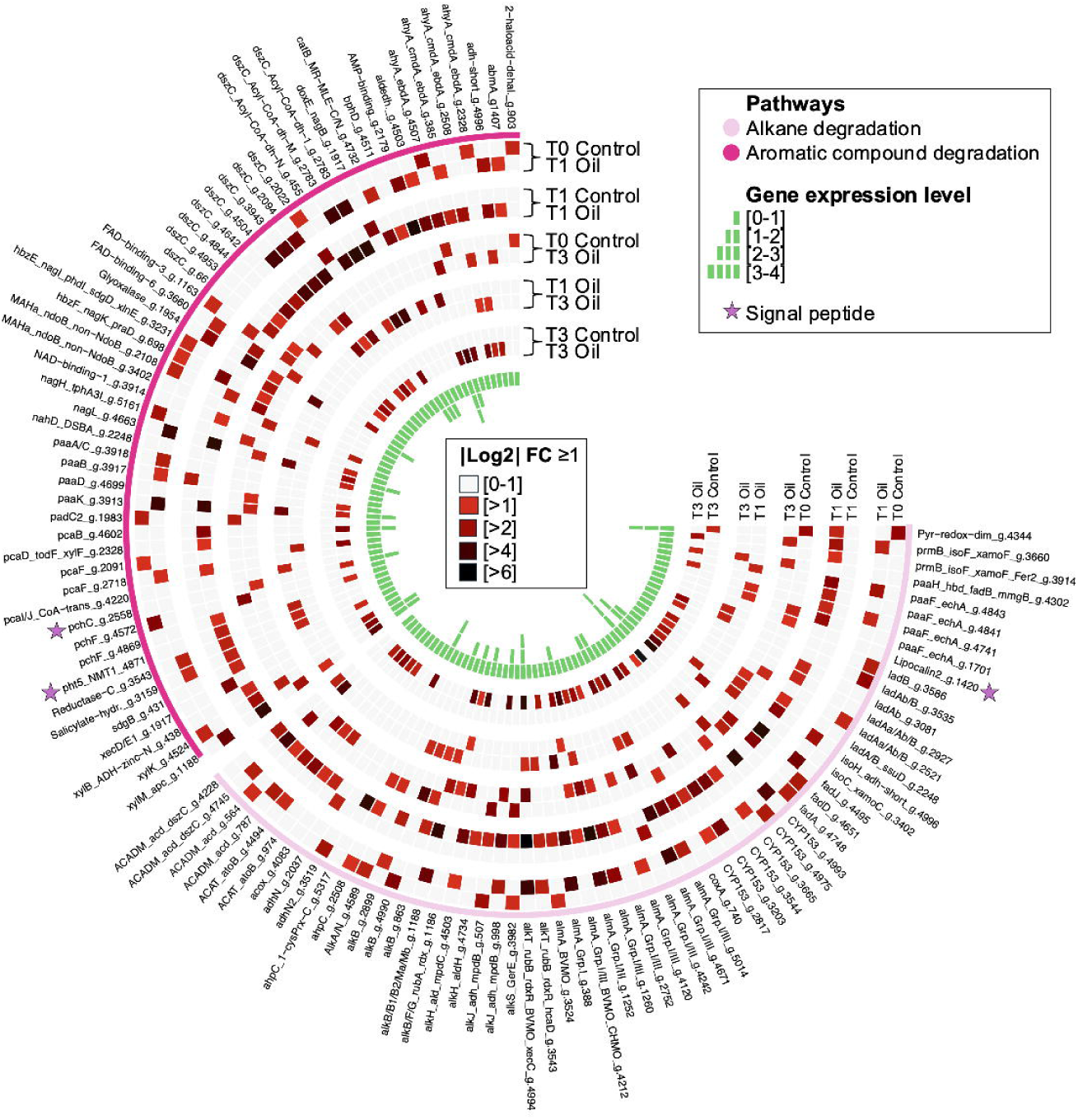

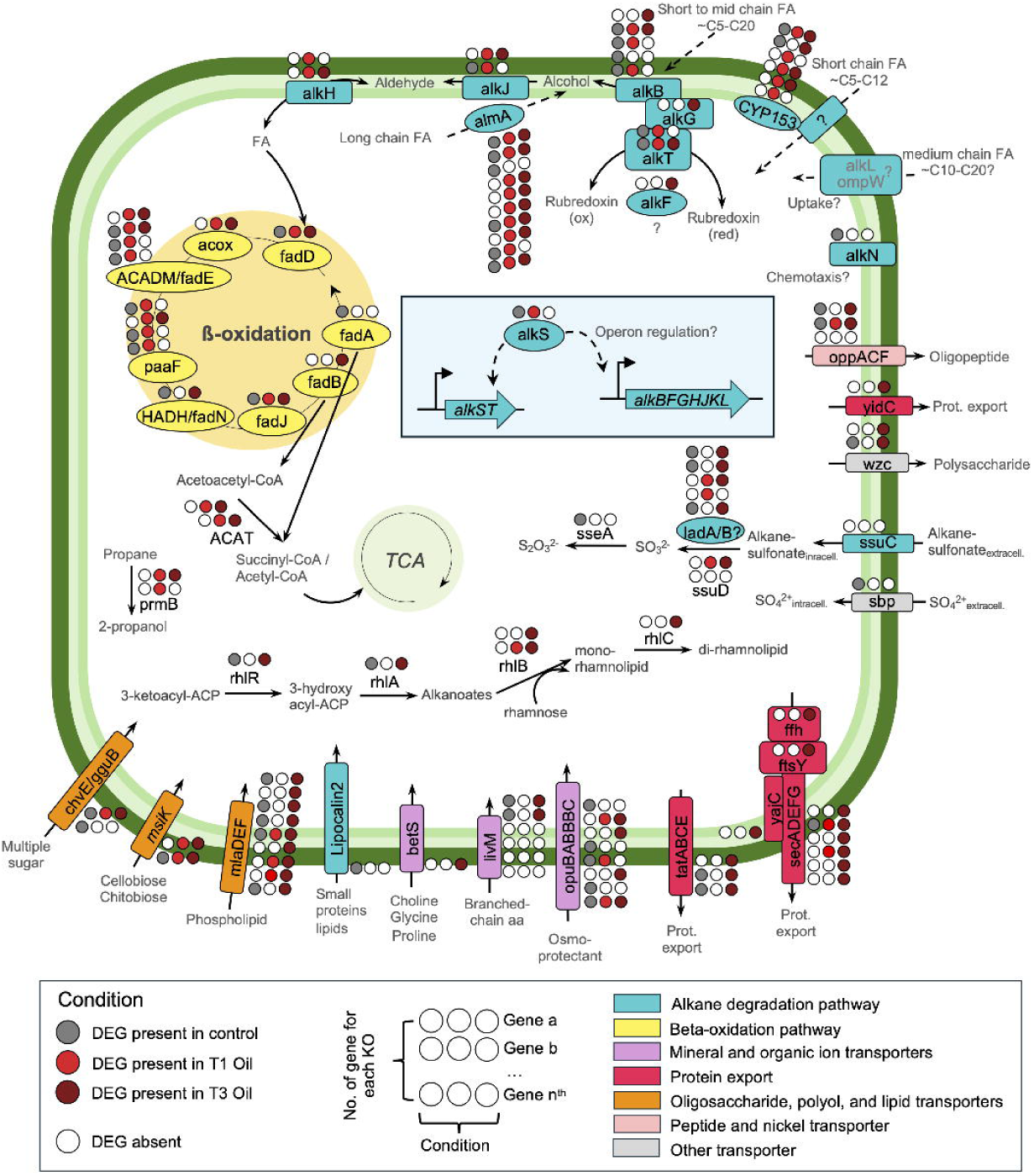

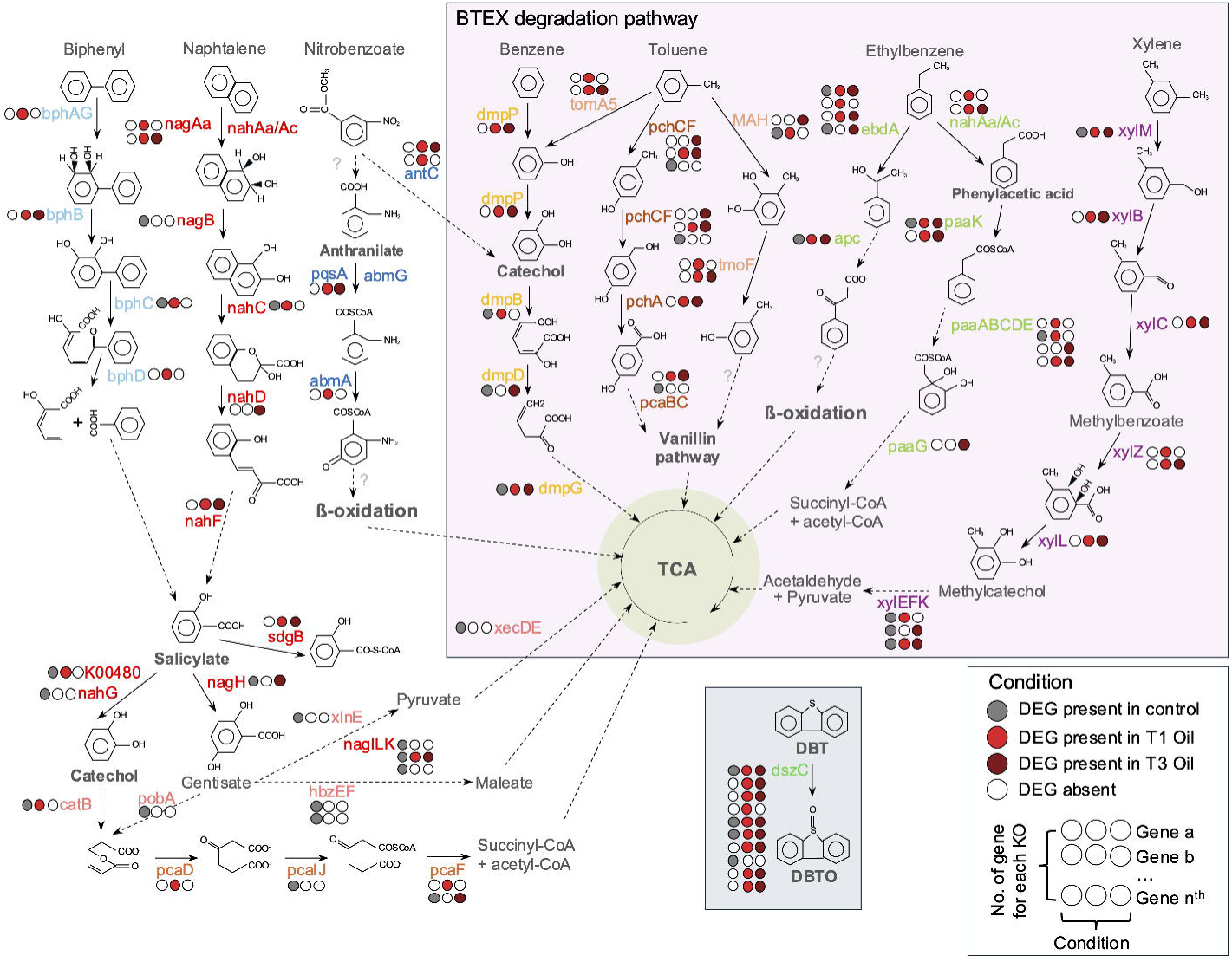

### Aliphatic and aromatic compounds degradation by *Rhodococcus*

A total of 61 and 55 DEGs were identified coding for proteins involved in alkane and aromatic compound degradation pathways, respectively (**Figs. 6-8, Tables S11-S13**). The most frequently detected protein involved in the alkane degradation pathway was the long-chain alkane oxidizing enzyme, *almA* group I and III, with or without the presence of the pfam domains BVMO (PF00743, PF07992 and PF13450) and/or CHMO (PF00743; **Fig. 7, Table S11**). Of the ten DEGs coding for *almA*, some showed consistent expression across all conditions. However, five out of ten DEGs coding for *almA* displayed selective expression, appearing exclusively in either control samples (without ULSFO, 1 DEG) or those exposed to ULFSO (4 DEGs). Several DEGs in the alkane degradation pathway, such as rubredoxin proteins 1 (*alkF*, PF00301) and 2 (*alkG*, PF00301), and aldehyde dehydrogenase (*alkH*, PF00171), were expressed only in ULSFO conditions at T1, T3, or both.

In the aromatic compound degradation pathways, *Rhodococcus* strain R1B_2T exhibits DEGs encoded essential enzymes for the degradation of BTEX, biphenyl, naphthalene, nitrobenzoate and dibenzothiophene (DBT; **Fig. 8, Table S11**). Notably, many DEGs coding for enzymes involved in the final steps of converting catechol and into smaller molecules that enter the TCA cycle, and the TCA cycle were expressed in the control condition. These enzymes include salicylate hydroxylase (K00480), salicylate hydrolase *nahG* (K00480) which converts salicylate to catechol, and salicylate 5-hydroxylase (*nagH*, PF13577) that converts salicylate to gentisate. Gentisate is further broken down into pyruvate via gentisate 1,2-dioxygenase (*nagI*, Uniprot O86041), fumarylpyruvate hydrolase (*nagL*, O86042), maleylpyruvate isomerase (*nagK*, O86043), and *xlnE* (Q9S3U6), or to maleate via *hbzEF* (PF07883 and PF01557). In contrast, some DEGs were only expressed in response to ULSFO exposure. For instance, in the xylene degradation pathway, enzymes such as toluene methyl-monooxygenase (*xylM*, K15757, PF00487), xylulokinase (*xylB*, K00854, PF00107, PF08240), benzaldehyde dehydrogenase (*xylC*, K00141), benzoate/toluate 1,2-dioxygenase reductase component (*xylZ*, K05784), and dihydroxycyclohexadiene carboxylate dehydrogenase (*xylL*, K05783) were mainly up-regulated at T1 and/or T3 incubated with ULSFO. Additionally, 10 DEGs coding for dibenzothiophene monooxygenase (*dszC*, K22219, PF02770, PF02771, PF08028), involved in converting DBT to DBT-sulfoxide (DBTO), were identified. Among the 10 DEGs coding for *dszC* varied expression patterns were displayed, with some being active in all conditions, while others were expressed exclusively in T1 and/or T3 under ULSFO exposure or control conditions.

### Carbohydrate active enzymes potentially involved in degradation

The analysis of carbohydrate-active enzymes (CAZyme) (Cantarel et al., 2009) revealed 40 distinct CAZy families, with glycoside hydrolases (GHs) comprising 45% of the total, glycosyltransferases (GTs) making up 30%, auxiliary activities (AAs) accounting for 15%, carbohydrate esterases (CEs) contributing 7.5%, and carbohydrate-binding modules (CBMs) representing 2.5%. Among the 74 DEGs identified as encoding CAZyme (**Fig. S6, Table S12**), 17 were found to have a signal peptide. This signal peptide, an extension of the amino acid sequence, indicates where the protein is destined to be transported, whether inside or outside the cell (Briggs and Gierasch, 1986). DEGs containing signal peptides were predominantly associated with the CE, CBM, and GH families. Several DEGs were exclusively expressed during the ULSFO exposure periods (1 and/or 3 months). These included 7 DEGs related to AA families (e.g., AA1, AA2, AA3, AA3_2, AA7), 2 DEGs encoding CBM48, 9 DEGs associated with CE families (e.g., CE1, CE5, CE14), 12 DEGs encoding GH families (e.g., GH3, GH13_3, GH13_10, GH13_11, GH13_20, GH15, GH43_23, GH77), and 15 DEGs related to GT families (e.g., GT2, GT2_3, GT4, GT20, GT35, GT39, GT51, GT87, GT89).

### Other key processes and genes

Further in-depth analyses using the HADEG database (Rojas-Vargas et al., 2023) identified DEGs involved in biosurfactant production, polymer degradation, cold resistance, protein export, secretion systems, osmoprotection, organic matter degradation, and methane cycling (**Fig. S7, Table S13**). Of the 78 DEGs, 8 contained signal peptides linked to osmoprotection, polymer degradation, and transporter functions. The final steps of rhamnolipid biosynthesis in *Rhodococcus* were exclusively expressed under ULSFO conditions (T1 and T3 Oil; **Figs. 7, S7**), involving helicase domain proteins (PF00270, PF00271 in *rhlB*; PF13641 in *rhlC*). Additionally, two DEGs expressed in control and T3 Oil encoded an HTH-type quorum-sensing regulator (*rhlR/gerE*, PF00196, PF03472) and a 3-(3-hydroxydecanoyloxy) decanoate synthase (*rhlA*, PF00561). Twenty DEGs related to polymer degradation were found, 11 of which were exclusive to ULSFO exposure. Two *cspA* DEGs (K03704) were upregulated in T3 Oil, and two *groEL* DEGs (K04077) were expressed in T3 Oil, with one also in the control condition.

For osmoprotection, 8 DEGs were identified in the osmoprotectant transporter system: 3 encoding the ATP-binding protein *opuBA* (K05847), 2 for the permease protein *opuBB* (K05846), and 2 with signal peptides for the substrate-binding protein *opuBC* (K05845). One DEG encoded *opuBC/AC* (K05845, PF04069) with a substrate-binding domain for the ABC-type glycine betaine transport system (**Figs. 7, S7**). Four DEGs (*opuBA* g.4676, *opuBB* g.4677, *opuBC* g.4674, g.4675) formed a cluster, exclusively expressed under ULSFO conditions. All 14 DEGs related to secretion and protein export were also expressed under ULSFO (**Figs. 7, S7**), including a membrane protein insertase *yidC* (K03217, PF02096), the preprotein translocase complex *yajC (*K03210), *secADEFG* (K03070, K03072, K12257, K03073, K03074 and K03075), and sec-independent translocases *tatA/E* (K03116), *tatB* (K03117), and *tatC* (K03118). Six DEGs encoding cytochrome p450, involved in transporting FA (C10-C20), were found, but no DEGs for short-chain FA transporters like *alkL* or *ompW* were detected.

## DISCUSSION

To adapt to external changes or environmental stressors, cells adjust their transcriptome to maintain internal stability (López-Maury et al., 2008). In the context of our study, we define stress as the exposure to ULSFO, which represents a substantial environmental challenge requiring metabolic adaptation for the bacterial cells. If such a perturbation persists, cells can acclimate to the new conditions through transcriptional adjustments and continue growing (Kültz, 2005). With the expected decrease in sea-ice conditions in the NWP, ice-influenced coastal Arctic environment will be exposed to potential oil spills from increased ship traffic. Therefore, we simulated a petroleum marine environment, exposing an Arctic coastal marine beach sediment isolate, *Rhodococcus* sp. strain R1B_2T, to ULSFO for three months. Our findings revealed multiple DEGs associated with hydrocarbon degradation (HD), including alkane and polycyclic aromatic HD, biosurfactant production, plastics/polymer degradation, osmoprotection, protein export, secretion systems, and various transporters.

### *Rhodococcus’* dynamic response to ULSFO exposure

We observed dynamic gene expression changes over time (T0, T1, T3) in *Rhodococcus* sp. strain R1B_2T in response to ULSFO exposure, with some pathways showing immediate and sustained upregulation, reflecting an urgent cellular response (Leahy and Colwell, 1990; Mohn and Tiedje, 1992). Petroleum hydrocarbon biodegradation analyses (**Fig. 1**) revealed different degradation patterns between compound classes: alkanes showed higher removal rates during the first month (T0-T1) with diminishing rates thereafter (T1-T3), while aromatic compounds exhibited the opposite pattern with higher removal rates between months T1-T3. This sequential degradation aligns with previous studies indicating that alkanes are preferentially biodegraded due to their simpler structure, while aromatic hydrocarbons require more complex metabolic pathways and are typically degraded later (Van Hamme et al., 2003; Bento et al., 2005; Coulon et al., 2005). The lower PAH content of ULSFO (Yang et al., 2023) explains the low initial naphthalene concentrations (**Fig. 1D**). The observed biodegradation pattern reveals a progressive response where aliphatic compounds were degraded primarily during the first month, while aromatic compounds showed minimal degradation initially but increased degradation between months 1 and 3, suggesting differential temporal effectiveness in the bacterial treatment of these hydrocarbon classes. Later-stage gene upregulation, including acyl-CoA dehydrogenase and FA desaturase (**Figs. 4, S4**), suggests later-stage adaptation or secondary response mechanisms to hydrocarbons (Leahy and Colwell, 1990; Mohn and Tiedje, 1992; Head et al., 2006). This pattern is similar to other oil-degrading bacteria like *Alcanivorax borkumensis* (Schneiker et al., 2006), *Cycloclasticus* sp. (Teira et al., 2007) and *Oleispira antarctica* (Kube et al., 2013), where initial shock and/or toxicity is followed by adaptation. Pathways for organic matter degradation and biosurfactant production (**Fig. S7**) increased at later time points, suggesting a time-dependent adaptation. KEGG analyses (**Fig. S5**) showed upregulation of aromatic compound degradation and FA metabolism, with a strong response to BTEX, naphthalene, and anthracene (**Fig. 8**), suggesting microbial communities efficiently mobilize to degrade complex aromatic hydrocarbons.

### Multiple HD pathway plasticity of Arctic NWP beach sediment *Rhodococcus*

The organism’s response to oil exposure is demonstrated by the activation of the *alkS* gene (**Fig. 7**), which may regulate the *alkST* and *alkBFGHJKL* operons responsible for alkane degradation (van Beilen et al., 2001). The expression of *alkS* was detected only under control conditions and after one month (T1) in the presence of ULSFO (**Figs. 6-7**). This suggests that while these operons are used during normal conditions, their genes are selectively expressed to produce alkane degradation proteins activated after one month. Previous studies have shown that *Rhodococcus* species possess multiple *alkB* operons (Whyte et al., 2002). Strain R1B_2T contains several genes related to alkane degradation pathways, including *alkB*, *ladA*, and *almA* (**Fig. 7**), highlighting its potential for effective application in oil-contaminated environments, especially under cold and saline conditions, since this strain is a halotolerant psychrophile (Lirette et al., 2024).

The upregulated genes highlighted in this study are specifically upregulated in response to hydrocarbons, reflecting a regulated expression mechanism triggered by environmental stimuli (here ULSFO). In *Rhodococcus*, this regulation likely serves as an adaptive strategy to conserve resources by limiting metabolic pathway expression to necessary survival conditions (Larkin et al., 2005). This selective expression highlights *Rhodococcus*’ evolved ability to activate specific pathways under certain conditions, enhancing its efficiency in bioremediation, where hydrocarbons signal the activation of genes for degrading complex oil components (de Carvalho, 2019). The role of environmental context in gene regulation and metabolic activity in *Rhodococcus* is crucial. Further studies on the signaling mechanisms involved and comparisons with other hydrocarbon-degrading microorganisms could offer valuable insights. Our phylogenetic analysis (**Fig. 2**), which included 54 known species, shows that our novel Arctic NWP beach sediment strain clusters apart from other polar *Rhodococcus* species like *R. erythropolis* and *R. qingshengii,* both known for their capacity to degrade hydrocarbons (Laczi et al., 2015; Lincoln et al., 2015), indicating a distinct evolutionary path and enhancing our understanding of the genus’ taxonomy and ecological roles.

While our genomic analyses confirmed that strain R1B_2T represents a subspecies of *R. cercidiphylli* rather than a novel species, important metabolic and functional differences distinguish it from other known Arctic hydrocarbon-degrading *Rhodococcus* species. Specifically, our pangenome analysis (**Fig. S1**) revealed that R1B_2T possesses several unique gene clusters not present in *R. erythropolis* and *R. qingshengii*. Furthermore, R1B_2T shows substantial genomic divergence from these known Arctic degraders, sharing only 76.27% and 76.24% ANI with *R. qingshengii* and *R. erythropolis* respectively (**Fig. 2B, Table S3**), compared to 98.9% with *R. cercidiphylli*. These genomic differences likely explain the strain’s distinctive temporal pattern of hydrocarbon degradation observed in our experiments, where adaptation to Arctic conditions has potentially resulted in specialized metabolic pathways optimized for the sequential degradation of different hydrocarbon classes in cold environments.

When comparing our strain’s hydrocarbon degradation strategy with other cold-adapted hydrocarbon degraders, several distinctive features emerge. While psychrophilic hydrocarbon degraders like *Oleispira antarctica* RB-8 shows specialization primarily in alkane degradation with limited aromatic degradation capabilities (Kube et al., 2013), and *Colwellia psychrerythraea* 34H has been characterized as a versatile cold-adapted heterotroph (Methé et al., 2005), our strain shows a more comprehensive degradation capacity across multiple hydrocarbon classes. Unlike many cold-adapted bacteria that typically show continuous expression of degradation genes, our strain exhibits precisely regulated temporal expression of *alkB* and related genes. Furthermore, while most characterized hydrocarbon degraders typically show relatively consistent degradation rates across exposure periods (Whyte et al., 2002), our strain demonstrates a distinctive sequential pattern— prioritizing alkanes initially before shifting to aromatic compounds. This strategic approach likely represents an evolutionary adaptation to maximize energy efficiency in cold, nutrient-limited environments, where conservation of cellular resources is crucial for survival.

Additionally, the synergistic relationship between *alkB, CYP153*, and *almA* pathways observed in our co-expression network (**Fig. 5D**) appears to be a unique feature not previously reported in other cold-adapted *Rhodococcus* strains such as *R. qingshengii* (Lincoln et al., 2015) and *R. erythropolis* (Laczi et al., 2015), suggesting our strain has evolved specialized regulatory mechanisms for coordinating multiple degradation pathways in response to complex hydrocarbon mixtures.

### Energy regulation in *Rhodococcus* exposed to ULSFO

We observed prominent features of the cytoplasm, cytoplasmic vesicles, membrane and extracellular regions components (**Fig. 4**), indicating shifts in energy metabolism. These components are vital for maintaining cellular integrity and function during hydrocarbon exposure (Harms and Bosma, 1997; Margesin and Schinner, 2001; Van Hamme et al., 2003). An increased expression of several dehydrogenases was observed, such as acyl-CoA and carbon monoxide dehydrogenase in *Rhodococcus* strain R1B_2T (**Figs. 4, S4-S5**), which suggests shifts in energy production pathways essential for managing oil-induced stress (Lindquist and Craig, 1988; Margesin and Schinner, 2001). The up-regulation of genes encoding acyl-CoA dehydrogenase, indicating degradation of aromatic compounds, and the increased FA and alkane metabolism, suggest active degradation of aromatic compounds (**Figs. 4, S4-S5**). Furthermore, many degradation pathways converge on the TCA cycle (**Figs. 7-8**), highlighting its central role in energy production, and the upregulation of the TCA cycle and oxidative phosphorylation pathways (**Figs. 4, S4**), supporting the notion of elevated energy production in strain R1B_2T, crucial for the energy-demanding process of HD (Atlas, 1981; Margesin and Schinner, 2001). Moreover, changes in nitrogen and sulfur cycling pathways indicate broad metabolic shifts (**Figs. S5, S7**), likely associated with the efficient processing and detoxification of oil components (Zhu et al., 2001; Knapik et al., 2020). The regulation of genes involved in methane metabolism and organic matter degradation points to the organism’s adaptive metabolism, utilizing oil components as substrates (Toth and Gieg, 2018). Additionally, we observed increased activity in FA biosynthesis, elongation, and metabolism pathways in oil-exposed samples (**Fig. S5**), which indicates an adaptation in lipid metabolism in response to the presence of oil, likely as part of the cellular response to utilize aliphatic compounds as a carbon sources (Leahy and Colwell, 1990; Mohn and Tiedje, 1992).

### Possible role of key CAZymes in hydrocarbon degradation (HD)

CAZymes play an important role in HD, reflecting complex adaptive responses in oil-contaminated environments. Various CAZyme families showed upregulation in oil-exposed conditions (**Fig. S6**), indicating their involvement in hydrocarbon metabolism. The AA family enzymes AA2 (catalase-peroxidase) and AA1 (laccase-like oxidoreductase) contribute to managing reactive oxygen species (Levasseur et al., 2013; Alharbi et al., 2019) and oxidizing aromatic substrates (Scott Conor et al., 2023), respectively. AA3 enzymes aid in oxidizing aromatic and aliphatic polyunsaturated alcohols (Sützl et al., 2018), while CBM48 facilitates complex carbohydrate degradation by bringing substrates closer to enzymes (Salam, 2018). CE family enzymes, particularly CE1, and GH family enzymes (e.g., GH3, GH13, GH15, GH43, and GH74) are implicated in lignin degradation and benzoic acid metabolism, suggesting a role in breaking down complex hydrocarbons like PAHs (Dou et al., 2021).

Several bacterial strains express CAZymes to enhance oil biodegradation. For instance, *Pseudomonas putida* and *P. aeruginosa* produce various GH and CE enzymes crucial for breaking down complex carbohydrates and hydrocarbons (Margesin and Schinner, 2001). *Alcanivorax borkumensis* is vital for marine oil spill cleanup due to its alkane degradation and biosurfactant production (Yakimov et al., 1998). *Rhodococcus erythropolis* and *R. opacus* possess CAZy genes that enhance hydrocarbon breakdown (Larkin et al., 2005), while *Marinobacter* species contribute significantly to oil-degrading communities in marine oil spills (Gauthier et al., 1992). *Acinetobacter baumannii* (Towner, 1992) and *Bacillus subtilis* (Nimrat et al., 2019) also produce various relevant CAZymes. Emerging research highlights *Dietzia*, formerly *Rhodococcus* (Rainey et al., 1995), as significant in alkane degradation and biosurfactant production (Soltanighias et al., 2019).

The upregulation of CAZyme genes in strain R1B_2T under oil exposure (**Fig. S6**), particularly at later time points (T3), suggests their role in microbial adaptation, allowing *Rhodococcus* to first degrade aliphatic compounds before aromatics and PAHs. The observed faster degradation rates of alkanes (**Fig. 1**), supports that lower molecular weight hydrocarbons are more susceptible to biodegradation (Atlas, 1981; Coulon et al., 2005). This adaptation may involve alternative energy sources, cell wall modifications, or biosurfactant production to enhance oil degradation. Combining different enzyme families, like GH and CE, could yield synergistic effects for more efficient pollutant breakdown (Singh and Ward, 2004). However, further research is needed to confirm these enzyme activities and their specific roles in oil degradation.

### Activation of stress-response genes in Arctic NWP beach sediment *Rhodococcus*

The presence of stress-response proteins, such as chaperonin GroEL and SCP2 (**Figs. S4, S7**), in strain R1B_2T indicates mechanisms to mitigate oil exposure effects, highlighting the complexity of microbial responses (Singh et al., 2014). Stress-related proteins, including cold shock proteins CspA were upregulated at T3 Oil (**Fig. S7**), suggesting a stress response to oil exposure (Choudhary et al., 2024). Upregulation of two-component systems and type II polyketide biosynthesis in oil conditions (**Figs. 7, S5**) points to secondary metabolites involved in environmental sensing and stress adaptation, aiding survival (Stock et al., 2000; Singh et al., 2014) and competition under oil exposure (Head et al., 2006; Jeanjean et al., 2008). Additionally, the upregulation of GT enzymes for cell wall biosynthesis and pathways for glycerolipid and glycerophospholipid metabolism (**Figs. S4, S6**) indicates that membrane modification is occurring and is crucial for maintaining cellular integrity in response to oil stress. This suggests that the microorganisms are adapting their cell walls to harsh oil conditions (Harms and Bosma, 1997; Nikaido, 2003; Mazzella et al., 2005). Furthermore, the presence of osmoprotectant transporters (**Figs. 7, S7**) indicates adaptation to osmotic stress often found in oil-contaminated environments (Park and Park, 2018).

### Cellular transport systems and synergistic hydrocarbon degradation mechanisms

The upregulation of ABC-type transporters, particularly those involved in multidrug transport and cytochrome biosynthesis (**Figs. 7, S7**), suggests activation of detoxification mechanisms in response to oil exposure (Van Hamme et al., 2003; Nikaido and Pagès, 2012). Increased activity in these transport systems, especially ABC transporters, implies enhanced molecular transport processes essential for the uptake and metabolism of hydrocarbons and other compounds (Nikolopoulou and Kalogerakis, 2008; Brooijmans et al., 2009). Although the specific transporter responsible for the uptake of short- and long-chain FAs is not yet identified, we did observe the expression of DEGs encoding cytochrome p450 (*CYP153*), also known as P450 alkane hydroxylase that oxidizes mid-chain-length alkanes (∼C10 to C20; **Fig. 7**) (van Beilen et al., 2006; Fenibo et al., 2023; Suzuki et al., 2025). However, no DEGs were found for transporters such as *alkL* or *ompW*, which are involved in transporting FAs with carbon chains longer than C10 (Julsing et al., 2012). CYP153 is thought to cooperate with *alkB* in degrading various *n-alkanes* (Nikaido, 2003; Liang et al., 2016). Our co-expression network analysis revealed a functional relationship between key hydrocarbon degradation genes, particularly highlighting the synergistic mechanisms between *alkB*, *CYP153* and *almA* (**Fig. 5D**). This synergy likely represents a complementary degradation strategy where *alkB* preferentially oxidizes short to medium-chain alkanes (∼C5-C20) and >C30 with other functional enzymes (Fenibo et al., 2023; Suzuki et al., 2025), while CYP153 more efficiently targets shorter chains (∼C5-C12) with some substrate range overlap (van Beilen et al., 2006). Their shared connections with alcohol and aldehyde dehydrogenases in our network (group of node A; **Fig. 5D**, **Table S10**) illustrate the complete metabolic pathway where alkanes are sequentially oxidized to alcohols, aldehydes, and finally fatty acids entering central metabolism. As for flavin-binding alkane hydroxylase *almA,* and flavin-dependent alkane monooxygenase *ladA*, both oxidize long chain alkanes, C28-C36 and C10-C30, respectively (Xiang et al., 2023; Suzuki et al., 2025), indicating a comprehensive degradation capacity across the full spectrum of alkane chain lengths.

The various upregulated transport systems in *Rhodococcus* appear to be essential for nutrient uptake and maintaining cellular balance, facilitating the transport of oligopeptides, proteins, polysaccharides, and ions across cell membranes (**Figs. 7, S7**). The upregulation of genes related to protein export and secretion suggests increased extracellular activity, potentially linked to oil degradation or cellular defence mechanisms. Changes in transporter and protein export gene expression indicate modifications in cellular import/export functions in response to oil stress. The presence of signal peptides in many upregulated genes, which are crucial for protein targeting and secretion (Briggs and Gierasch, 1986), highlights the significance of extracellular enzymatic processes in adapting to oil contamination (Freudl, 2018).

### Importance of biosurfactant production in oil biodegradation

The synthesis of biosurfactants plays a crucial role in microbial adaptation to oil-contaminated environments. Our study identified 14 DEGs related to biosurfactant production, notably upregulating acyltransferase proteins and LuxR family regulators (like *rhlR*) after petroleum exposure (**Figs. 7, S7, Table S13**). LuxR regulators are essential for biosurfactant synthesis (Wei et al., 2004; Kothari and Jobanputra, 2022). Biosurfactants enhance the emulsification and solubilization of hydrophobic compounds, improving their bioavailability and degradation rates (Desai and Banat, 1997; Mulligan, 2005). Their amphiphilic properties facilitate oil droplet breakdown and increase the surface area for enzymatic degradation (Banat et al., 2010), thereby enhancing hydrocarbon uptake and metabolism in bacteria. In *Rhodococcus* strain R1B_2T, biosurfactant production, such as rhamnolipids, likely helps reduce the size of large oil droplets, increasing surface area for enzymatic action. This emulsification improves the bioavailability of hydrocarbons, allowing for more efficient uptake and metabolism (Ron and Rosenberg, 2002). The variety of biosurfactants, like rhamnolipids and surfactin, enables bacteria to adapt to different oils and conditions (Mulligan, 2005). Biosurfactants, like rhamnolipids, are well-known in hydrocarbon biodegraders, especially in high G+C content bacteria, like *Rhodococcus* species (Whyte et al., 1997; Raymond-Bouchard et al., 2018). Their biodegradable nature makes them environmentally favorable while persisting long enough to support oil degradation effectively. Previous studies on *Rhodococcus* species, including *R. erythropolis* (Pacheco et al., 2010) and *Rhodococcus* sp. strain TA6 (Pal et al., 2009), support these findings, despite a distant evolutionary relationship with strain R1B_2T (**Fig. 2**). Biosurfactant production is often upregulated in bacteria exposed to hydrophobic substrates, highlighting their role in natural degradation processes (Maier and Soberon-Chavez, 2000). Enhancing biosurfactant production in oil-degrading bacteria is crucial for improving bioremediation technologies. Additionally, strain R1B_2T shows upregulation of alkane hydroxylase genes (e.g., *alkB* and CYP153) in response to hydrocarbon exposure (**Figs. 6-7**), demonstrating metabolic flexibility in cold marine environments. Arctic *Rhodococcus* species’ ability to form biofilms and produce biosurfactants enhances their hydrocarbon degradation efficiency (Perfumo et al., 2010). These traits, particularly cold tolerance, make *Rhodococcus* a key player in developing bioremediation strategies for oil spill mitigation in Arctic coastal sediment and marine ecosystems.

## CONCLUSION

Our study reveals key novel insights into cold-adapted hydrocarbon degradation mechanisms. Unlike mesophilic *Rhodococcus* strains, our Arctic isolate demonstrates a distinctive sequential degradation strategy—preferentially metabolizing alkanes during early exposure before shifting to aromatic compounds later. This temporal orchestration represents a specialized adaptation to conserve energy in cold, oligotrophic conditions. Our transcriptomic analysis identified precisely timed expression patterns not documented in temperate strains, including the strategic expression of *alkS* only at specific timepoints. The co-expression network analysis revealed novel functional relationships between *alkB*, *CYP153*, and *almA* pathways that differ from the independent operation typically observed in mesophilic strains. When compared with other psychrotolerant hydrocarbon degraders, our *Rhodococcus* strain exhibits a more versatile metabolic toolkit for degrading multiple hydrocarbon classes, suggesting an evolved adaptation to the variable hydrocarbon mixtures in coastal Arctic environments. These findings advance our understanding of cold-adapted bioremediation mechanisms and provide valuable insights for developing Arctic-specific remediation technologies for the increasingly vulnerable Northwest Passage.

## SUPPLEMENTARY MATERIAL

The Supplementary Material for this article can be found online.

## ACKNOWLEDGEMENTS

We thank the community of Resolute Bay and in particular Devon Malik for providing logistical support, and his assistance in the field as our guide and bear watcher. We thank Ya-Jou Chen for her help in collecting samples in the field. We thank Canadian Polar Continental Shelf Program for providing logistical support for the field work.

## DATA AVAILABILITY STATEMENT

The raw transcriptome sequencing datasets generated in this study are available at NCBI under BioProject ID PRJNA1193011. Whole genome sequencing is available on JGI genome portal under Gold Sequencing Project ID Go0632451, Gold Analysis Project ID Ga0557218 and raw sequencing read are available at NCBI under BioProject ID PRJNA945214.

## AUTHOR CONTRIBUTIONS

**Nastasia J. Freyria**: Data curation; Formal analysis; Investigation; Methodology; Project administration; Software; Validation; Visualization; Writing – Original Draft Preparation; Writing – Review & Editing. **Antoine-Olivier Lirette**: Conceptualization; Methodology; Writing – Review & Editing. **Brady R. W. O’Connor**: Data curation; Methodology; Writing – Review & Editing. **Charles W. Greer**: Resources; Supervision; Writing – Review & Editing. **Lyle G. Whyte**: Conceptualization; Funding Acquisition; Project administration; Resources; Supervision; Validation; Writing – Review & Editing.

