## Supplementary material for "Transcriptomic analyses unveil hydrocarbon degradation mechanisms in a novel polar *Rhodococcus* sp. strain R1B_2T from a high Arctic intertidal zone exposed to ultra-low sulfur fuel oil": Fig. S1; Fig. S2; Fig. S3; Fig. S4; Fig. S5; Fig. S6; Fig. S7

**Table S1 (xlsx).** Petroleum hydrocarbon initial concentration, concentration measured, concentration of degradation and percentage of hydrocarbon removal from **Fig. 1.**

**Table S2 (xlsx).** Overall summary of results of the Illumina transcriptome sequencing.

**Table S3 (xlsx).** Genome information and average nucleotide identity values between selected reference genomes of genus *Rhodococcus* from the pangenome comparison from **Figs. 2 and S1.**

**Table S4 (xlsx).** Blastp results from comparison of *almA* gene annotated as putative flavin-binding monooxygenase from *Alloalcanivorax* *dieselolei* (ADP30851.1) with 8 referenced genomes of *Rhodococcus* species and novel strain R1B_2T.

**Table S5 (xlsx).** Statistic one-way ANOVA for total petroleum hydrocarbon analyses from **Fig. 1.**

**Table S6 (xlsx).** List of genes and functional annotation from clusters of top 25 heatmap from **Fig. 3.**

**Table S7 (xlsx).** List of Gene Ontology (GO) terms from **Fig. 4.**

**Table S8 (xlsx).** List of Clusters of Orthologous Genes (COG) terms from **Fig. S4.**

**Table S9 (xlsx).** List of KEGG eukaryotic Ortholog Groups of proteins (KO) terms from **Fig. S5.**

**Table S10 (xlsx).** List of genes present in each module and group of nodes from **Fig. 5D.**

**Table S11 (xlsx).** List of hydrocarbon degradation genes from **Figs. 6-8.**

**Table S12 (xlsx).** List of genes coding for CAZY enzymes from **Fig. S6.**

**Table S13 (xlsx).** List of genes from additional categories from **Figs. 7 and S7.**

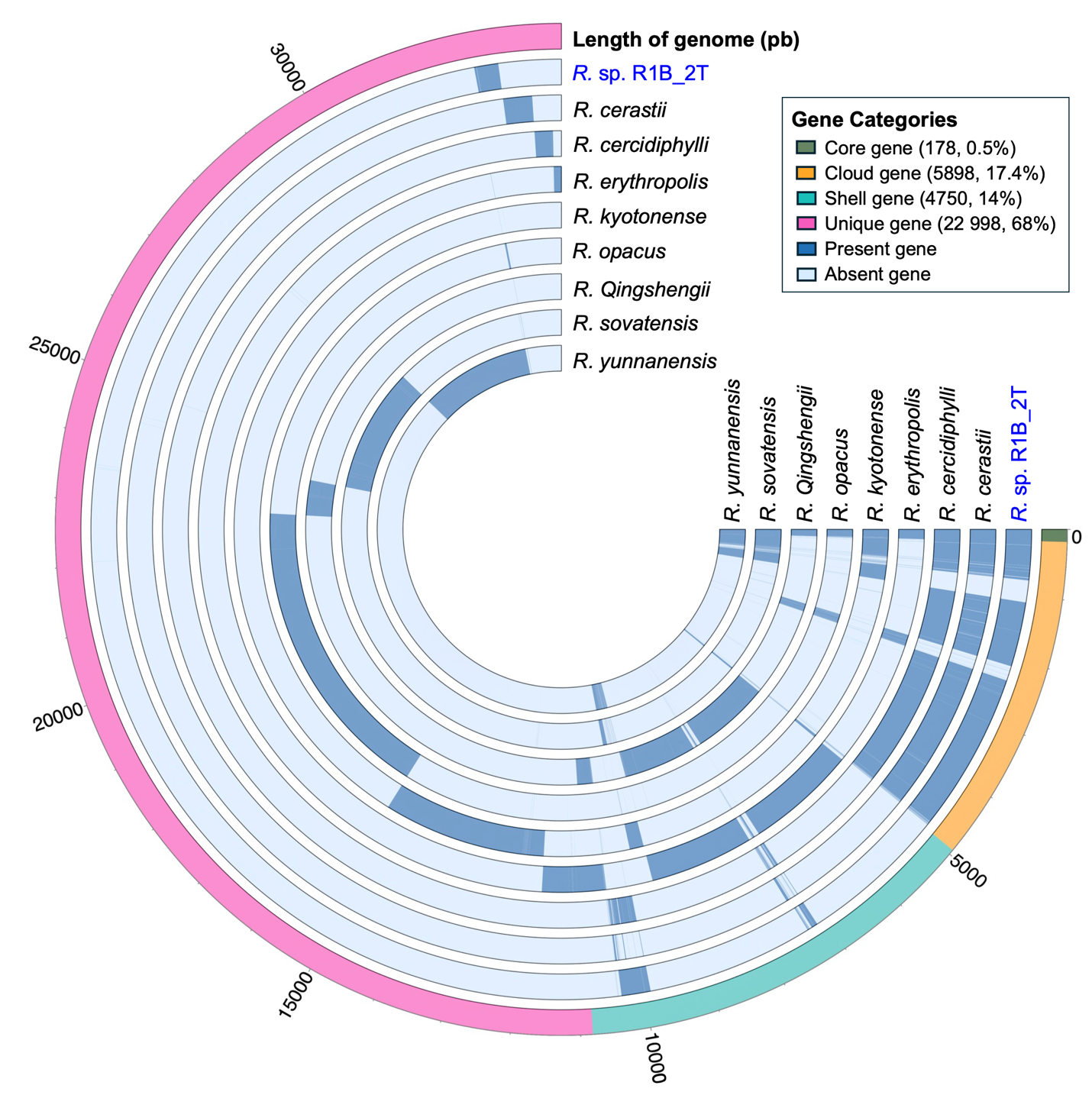

**Fig. S1. Pangenome of 10 *Rhodococcus* species**. The circular pangenome compares 9 genomes of referenced species of *Rhodococcus* with *Rhodococcus.* sp. strain R1B_2T. Core gene: genes present in all genomes; cloud gene: genes present in less than 15% of genomes; shell gene: genes present in 15-95% of genomes; and unique gene: genes present in only 1 genome. Genomes information can be found in **Table S3**.

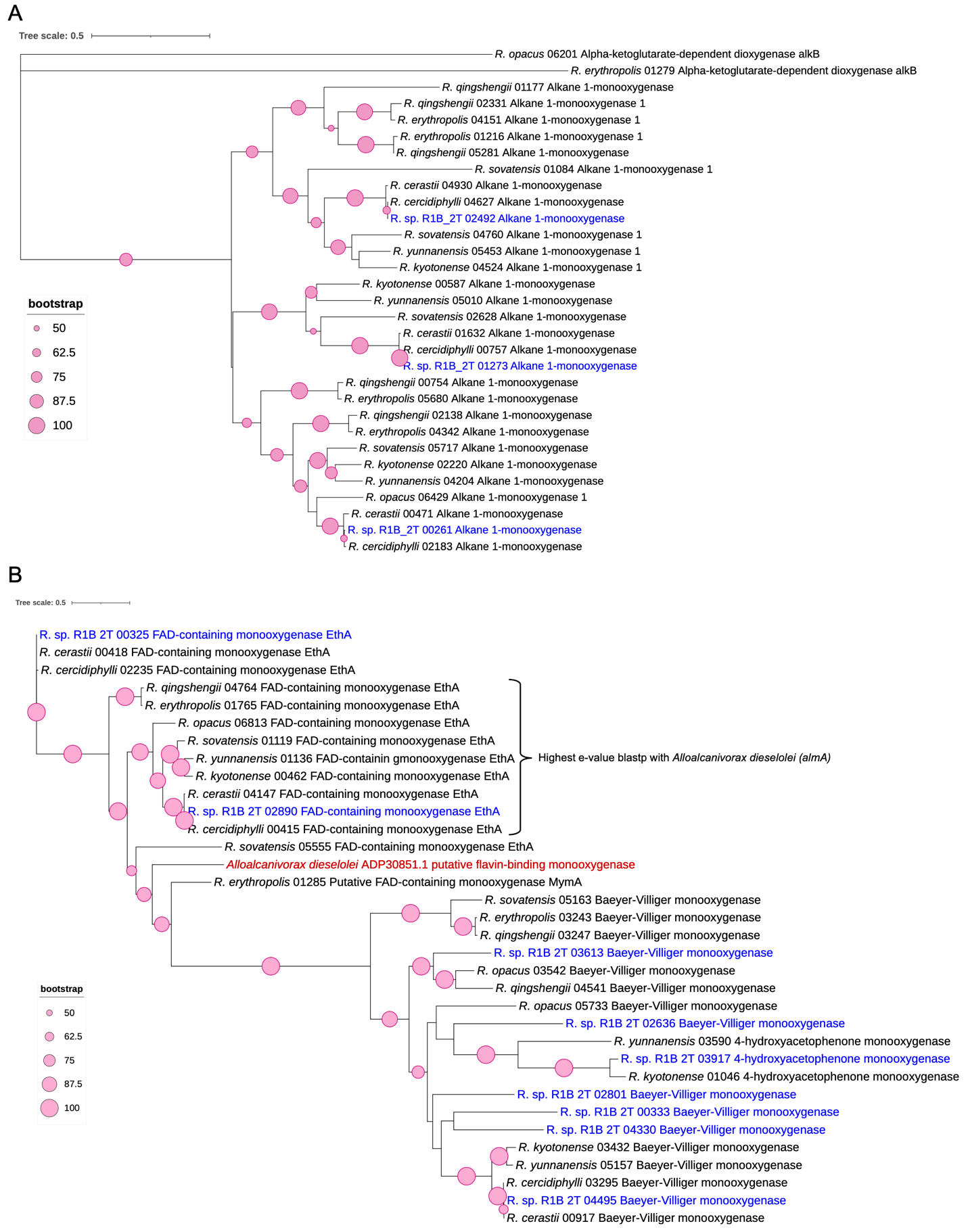

**Fig. S2. A Maximum Likelihood *alkB* and *almA* genes trees generated using RAxML**. (**A**) *alkB* genes phylogenetic tree constructed with 31 nucleic acid sequences of 1390 bp aligned along with 8 genomes of different species of *Rhodococcus* and with novel strain R1B_2T. (**B**) *almA* genes phylogenetic tree constructed with 34 amino acid sequences of 560 aa aligned along with 8 genomes of different species of *Rhodococcus* and with novel strain R1B_2T. The trees were constructed with bootstrap support calculated over 1,000 repetitions. Only bootstrap values greater than 50 (out of 100) are displayed. Blastp results for *alma* genes can be found in **Table S4**.

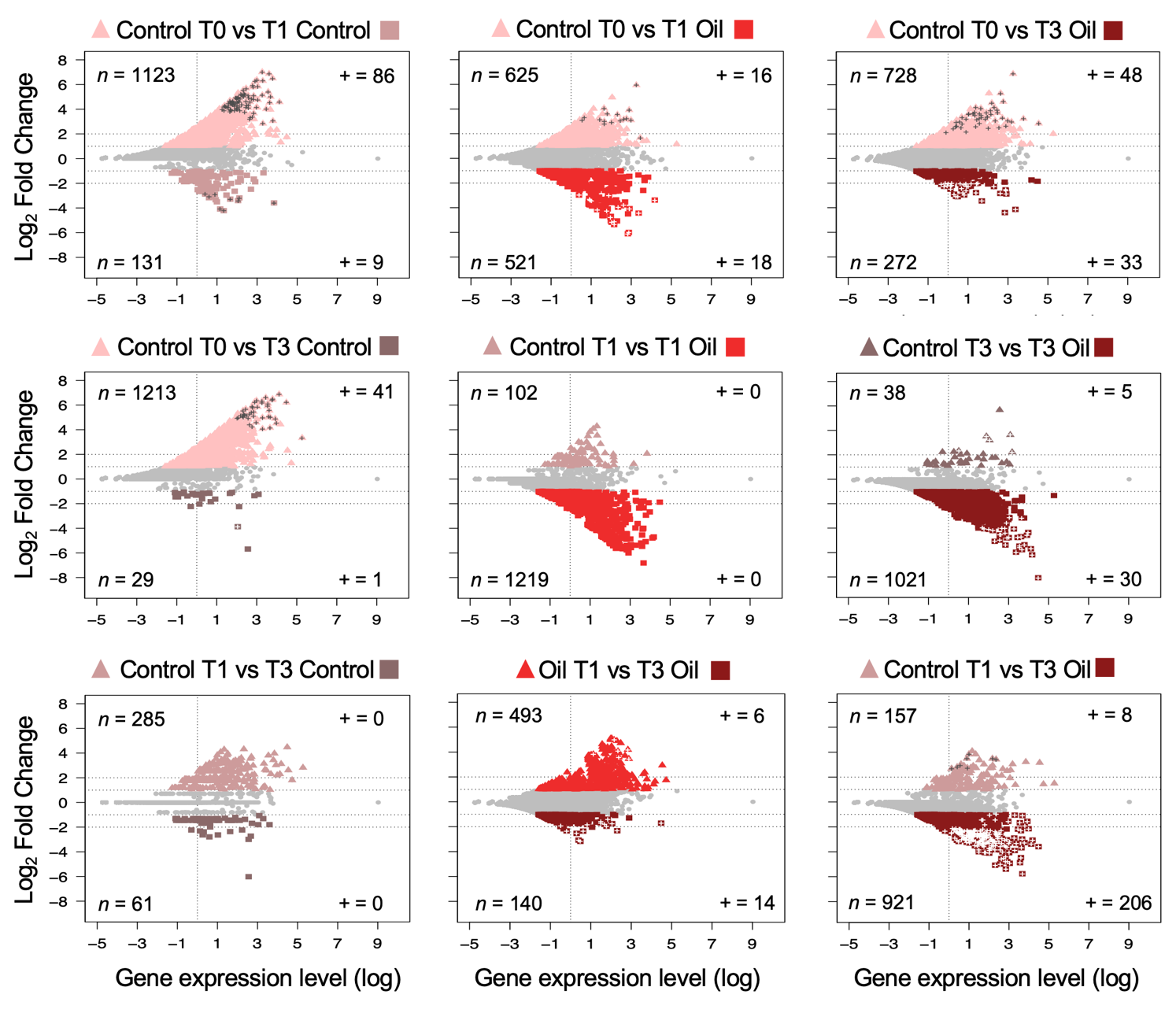

**Fig. S3. Differential expression genes (DEGs) between time of sampling (T0, T1 and T3) with and without oil.** Scatter plots show the comparison between months (T), T1 – 1 month; T3 – 3 months) of cells exposed either with ultra-low sulfur fuel oil (ULSFO) or without oil (control). Each point represents a unigene. Points greater than 1 of log_2_ Fold Change (FC) indicate up-regulated genes and lower than -1 indicate down-regulated genes. Color points indicate differential expressed genes (DEGs). Symbol “+” indicate only filtered DEGs with an Benjamini-Hochberg-adjusted *p*-value <0.05.

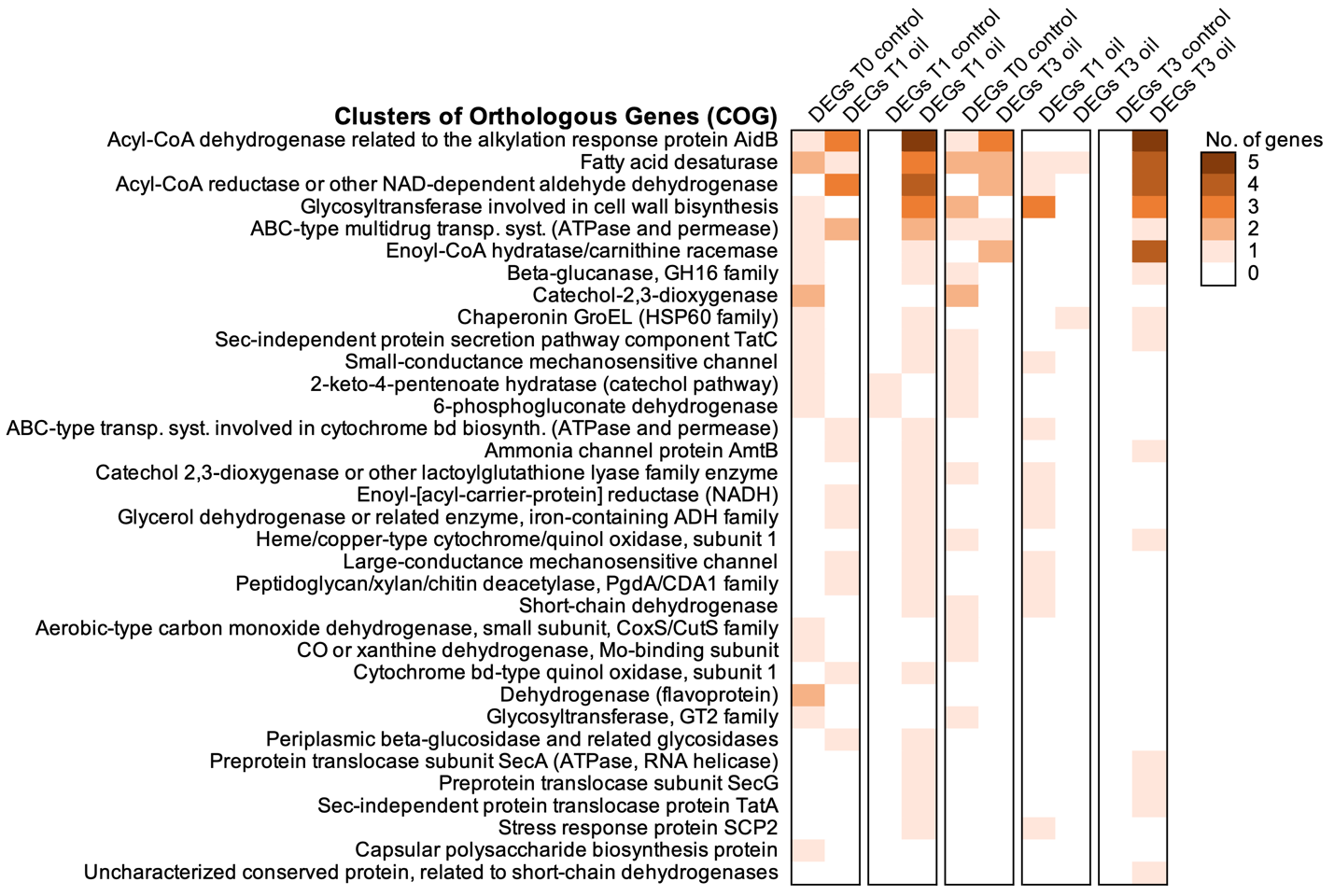

**Fig. S4. Top expression of differentially expressed genes (DEGs) annotated with Clusters of Orthologous Genes (COG) between comparisons of time and with or without the presence of ULSFO.** A complete list of genes and annotations can be found in **Table S8**.

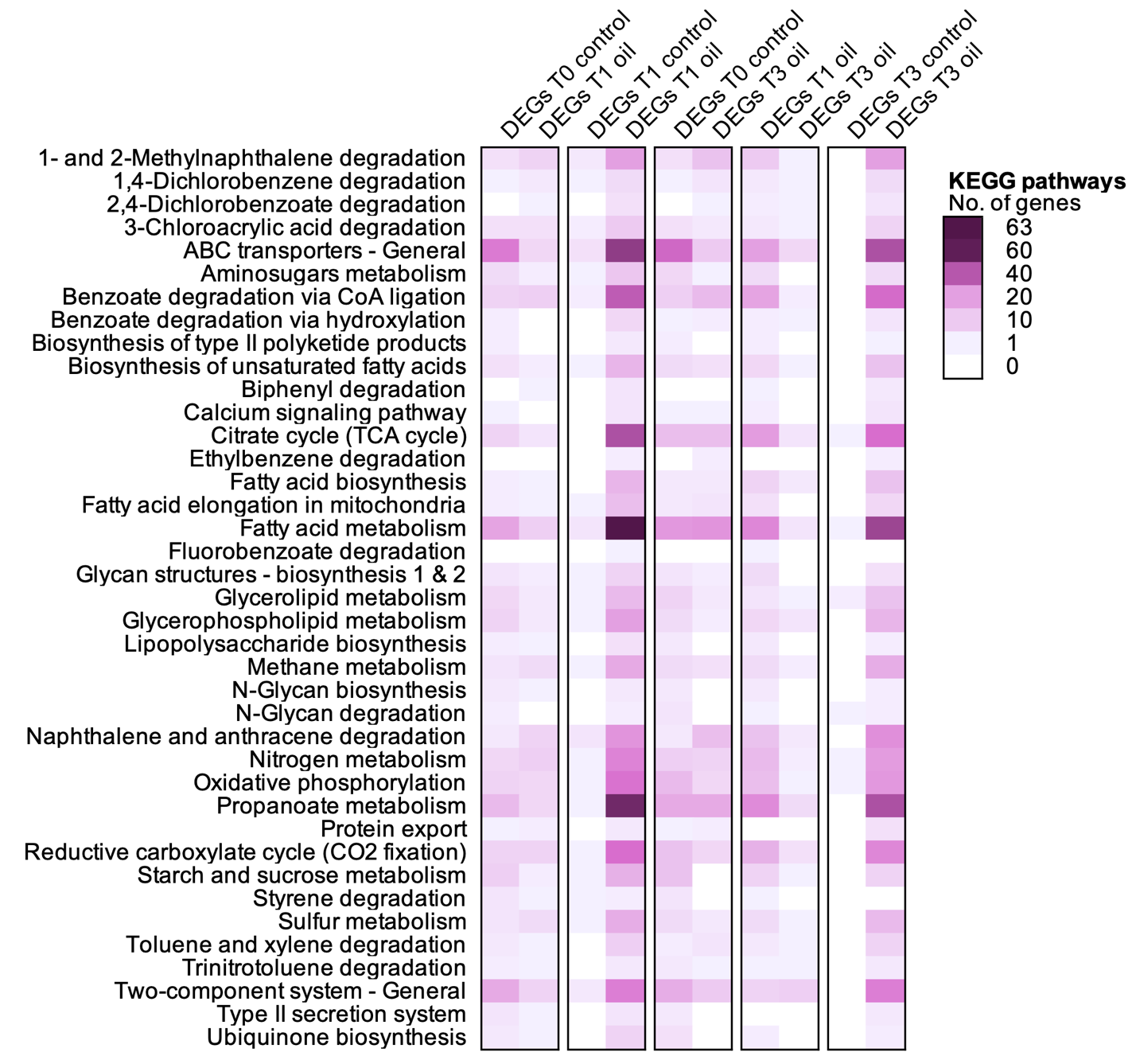

**Fig. S5. Top expression of differentially expressed genes (DEGs) annotated with KEGG Orthology (KO) between comparisons of time and with or without the presence of Ultra-Low Sulfur Fuel Oil (ULSFO).** A complete list of genes and annotations can be found in **Table S9**.

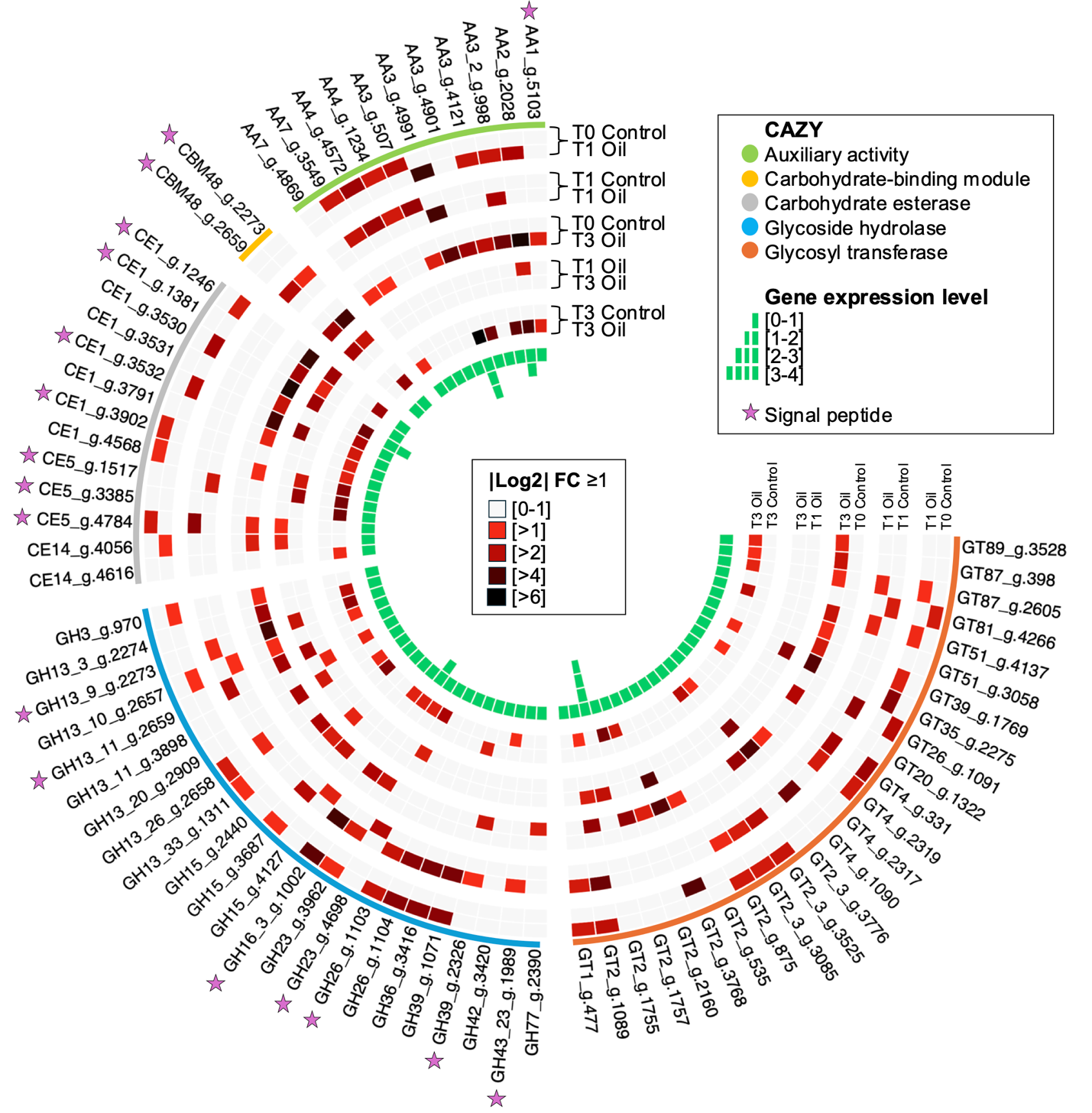

**Fig. S6. Circular heatmap of differentially expressed genes (DEGs) annotated as** **carbohydrate-active enzymes (CAZy) categories.** The heatmap depicts expression profiles based on log2 fold changes across sampling timepoints (T0, T1 - 1 month, T3 - 3 months) under conditions with and without Ultra-Low Sulfur Fuel Oil (ULSFO). The outer circle color-codes each DEG according to its specific CAZy category. Expression intensity is represented by color gradient, with darker shades indicating higher expression levels. **Table S12** provides a comprehensive list of all genes with their annotations.

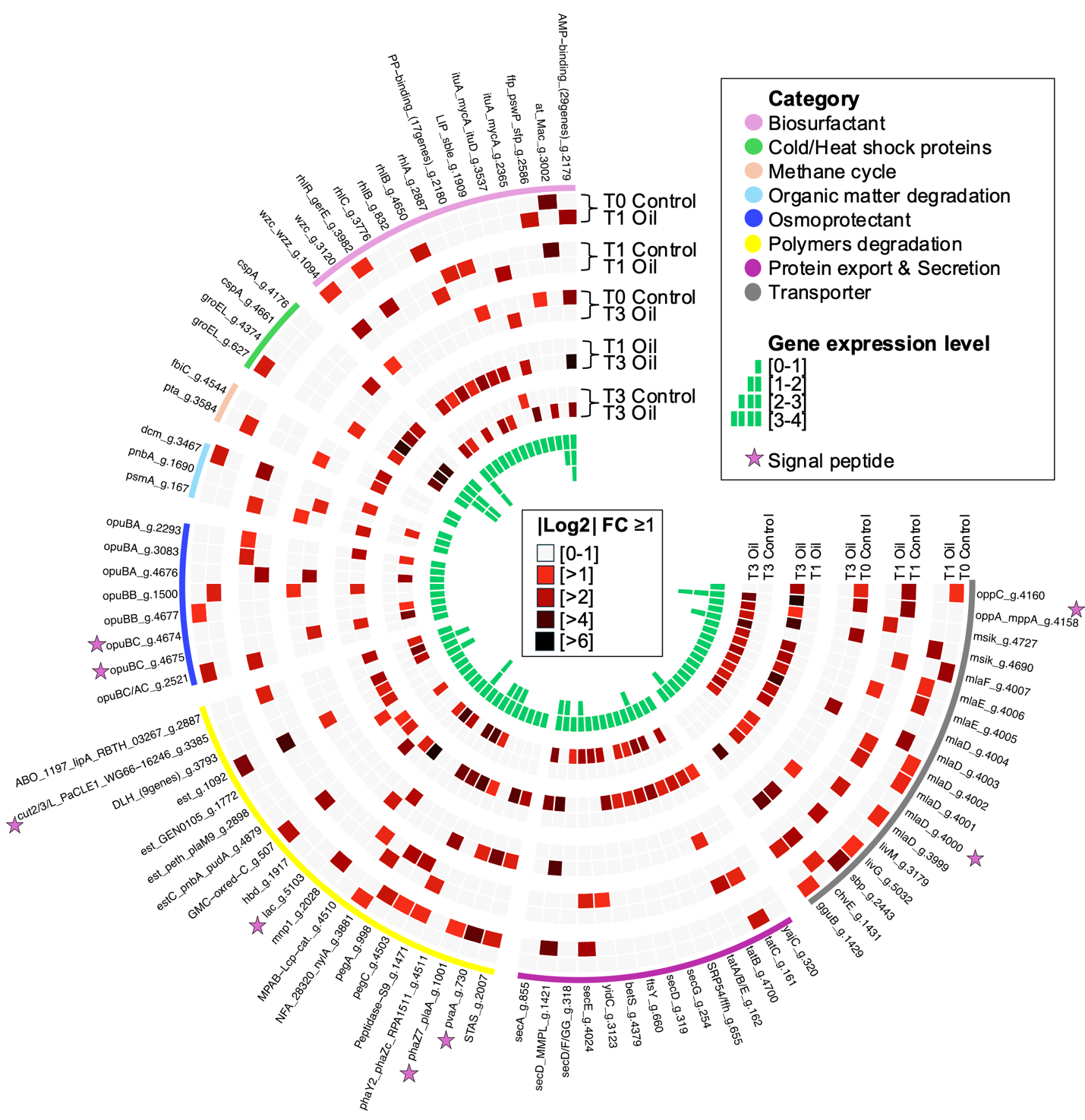

**Fig. S7. Circular heatmap of differentially expressed genes (DEGs) annotated as** **different pathway categories.** The heatmap depicts expression profiles based on log2 fold changes across sampling timepoints (T0, T1 - 1 month, T3 - 3 months) under conditions with and without Ultra-Low Sulfur Fuel Oil (ULSFO). The outer circle color-codes each DEG according to its specific pathway annotation category. Expression intensity is represented by color gradient, with darker shades indicating higher expression levels. **Table S13** provides a comprehensive list of all genes with their annotations.
